# A New Information Theoretic Approach Shows that Mixture Models Outperform Partition Models for Phylogenetic Analyses of Amino Acid Data

**DOI:** 10.64898/2026.03.16.712229

**Authors:** Huaiyan Ren, Changsen Jiang, Thomas K.F. Wong, Yutong Shao, Edward Susko, Bui Quang Minh, Robert Lanfear

**Affiliations:** School of Computing, College of Systems & Society, Australian National University, Canberra, ACT 2600, Australia; Ecology and Evolution, Research School of Biology, College of Science & Medicine, Australian National University, Canberra, ACT 2600, Australia; Mathematical Sciences Institute, College of Systems & Society, Australian National University, Canberra, ACT 2600, Australia; Department of Mathematics and Statistics, Faculty of Science, Dalhousie University, NS, Canada

**Keywords:** phylogenetics, model selection, partition model, mixture model, Akaike information criterion, parametric bootstrap

## Abstract

Partition and mixture models are widely employed in Maximum Likelihood phylogenetic analyses of large genomic datasets. Comparing the fit of the two types of models has been challenging, because standard information-theoretic approaches cannot be applied. Mixture models are increasingly popular for the analysis of amino acid datasets and can lead to different conclusions compared to partition models. This raises an important question – which type of model tends to perform better? Susko et al. (2026) recently introduced the marginal Akaike information criterion (mAIC), which allows mixture models and partition models to be directly compared for the first time. Here, we use the mAIC and a range of other approaches to compare the fit of mixture and partition models across a diverse set of empirical amino acid datasets. We show that the mAIC favours mixture models in all the datasets we examined. This has important implications for interpreting empirical analyses and suggests that continued development of mixture models is an important avenue for future research.

## Introduction

In phylogenetics, the use of large genomic sequence alignments has become increasingly common. It is well established that different sites evolve in different ways, and that accounting for this heterogeneity can be crucial for accurate phylogenetic inference (Cannon et al. 2016; Wang et al. 2019; Williams et al. 2021; Stiller et al. 2024; Giacomelli et al. 2025). To accommodate this heterogeneity, both partition models (Lanfear et al. 2017; Wang et al. 2019) and mixture models (Lartillot and Philippe 2004; Wang et al. 2008; Jayaswal et al. 2014; Ren et al. 2025) have been proposed, and one or both are implemented in many popular software programs (Guindon et al. 2010; Ronquist et al. 2012; Lartillot et al. 2013; Höhna et al. 2016; Bouckaert et al. 2019; Togkousidis et al. 2023; Wong et al. 2026).

In partition models, alignments are grouped a priori into distinct subsets, for example based on protein-coding genes, codon positions or other criteria (Nylander et al. 2004). A single substitution model is then assigned to each subset of sites. The alignment subsets with similar evolutionary models (e.g., similar relative substitution rates and/or equilibrium frequencies) can be merged using a variety of information criteria (e.g., Akaike information criterion (AIC), the corrected Akaike information criterion (AICc), and Bayesian information criterion (BIC)), thereby improving the overall partitioning scheme (Lanfear et al. 2012; Lanfear et al. 2017). The choice of partitioning scheme can be critical for partitioned analyses, as an inappropriate scheme has been shown to lead to incorrect phylogenetic inference in simulations (Wang et al. 2019) and has been shown to mislead empirical phylogenetic analyses (Brandley et al. 2005; Chiari et al. 2012; Kainer and Lanfear 2015). In addition, branch lengths may be inferred separately for each partition, depending on the partition model settings: partitions may share identical branch lengths (edge-equal), have branch lengths that differ but are proportionally scaled (edge-proportional), or have completely independent branch lengths (edge-unlinked, (Duchêne et al. 2020)).

Mixture models likewise estimate multiple evolutionary processes for a single alignment. They have been used to model heterogeneity in substitution processes (similar to edge-equal partition models), such as the EX3 and LG4X amino acid models (Le et al. 2008; Le et al. 2012) and DNA mixture models estimated by MixtureFinder (Ren et al. 2025). Mixture models have also been applied to capture the variation of evolutionary rates (similar to edge-proportional partition models, (Yang 1994; Soubrier et al. 2012)); the variation in branch lengths (similar to edge-unlinked partition models) in the GHOST model (Crotty et al. 2020); and heterogeneity in amino acid frequencies, as in the CAT profile models (Lartillot and Philippe 2004). In the original Bayesian formulation, the CAT model treats the equilibrium amino acid frequencies (profiles) as a Dirichlet process mixture, so that both the number of profiles and the profile parameters are estimated from the data (Lartillot and Philippe 2004). The C10–C60 models are a maximum-likelihood approximation to this, in which a fixed set of 10 to 60 profiles, pre-estimated from large empirical databases, is used together with a set of weights giving the prior probability of each profile and all profiles share a common exchangeability matrix (Quang et al. 2008). A key distinction from partition models is that mixture models do not assign sites to predefined subsets. Rather, in a mixture model the likelihood of each site is calculated under every model.

Both partition models and mixture models have been shown to improve phylogenetic inference in many empirical studies (Ruhfel et al. 2014; Cannon et al. 2016; Redmond and McLysaght 2021; Stiller et al. 2024; Giacomelli et al. 2025). However, a long-standing question in phylogenetics is whether one approach tends to perform better than the other. To evaluate and select among complex models, many studies have relied on simulation data to compare model performance. These comparisons have employed a range of criteria, including the accuracy of inferred tree topologies and branch lengths (Whelan and Halanych 2017), information criteria such as AIC or BIC (Jhwueng et al. 2014; Susko and Roger 2020; Liu et al. 2023), cross-validation (Susko and Roger 2020), bootstrap support (Jhwueng et al. 2014; Susko and Roger 2020; Liu et al. 2023), Kullback–Leibler divergence (Jhwueng et al. 2014; Liu et al. 2023) and posterior predictive simulation (Giacomelli et al. 2025). Most recently, Pisani et al. (2026) compared mixture, homogeneous and partition models in a large simulation study, assessing both the recovery of the true tree and model adequacy, and found that mixture models, and CAT-GTR in particular, outperformed the other two model classes. In addition, Redmond and McLysaght (2021) proposed a strategy that combines partition models and mixture models by applying mixture models (e.g., EX2, CAT) within partitioned alignments, further improving phylogenetic inference. However, most of these approaches, except AIC and BIC, are computationally expensive and are not implemented in user-friendly software programs or pipelines, which limits their application in empirical analyses. The upshot is that we still do not know whether, in general, mixture models or partition models tend to perform better for the phylogenetic analysis of empirical amino acid alignments.

Information criteria are relatively straightforward to calculate, as they rely solely on model parameters and likelihoods, and have been implemented in most phylogenetic software (Guindon et al. 2010; Ronquist et al. 2012; Lartillot et al. 2013; Höhna et al. 2016; Bouckaert et al. 2019; Togkousidis et al. 2023; Wong et al. 2026). Thus, they are among the most commonly used criteria for model selection, such as in ModelFinder (Kalyaanamoorthy et al. 2017), PartitionFinder (Lanfear et al. 2012) or MixtureFinder (Ren et al. 2025). However, existing information criteria have been shown to be inappropriate for comparing partition models with mixture models (Crotty and Holland 2022; Liu et al. 2023). Crotty and Holland (2022) simulated four-taxon alignments with two equal-size partitions and introduced incorrect partitioning by assigning 0–50% of sites to the incorrect partition. When partition models were fitted to these mispartitioned alignments, the AIC still tended to favour partition models over mixture models, even when 30% of the sites were incorrectly assigned and with large numbers of sites. Moreover, when the partitioning scheme was imperfect, the partition models (but not the mixture models) led to the inference of incorrect tree topologies.

Recent studies (Liu et al. 2023; Susko et al. 2026) suggest that this bias of model selection results from the different types of likelihood calculation in partition and mixture models. Specifically, Susko et al. (2026) pointed out that partition models compute likelihoods as conditional probabilities, since sites within each partition are assigned their own optimal substitution model. In contrast, mixture models calculate likelihoods as marginal probabilities, because sites are not associated with fixed partition labels. Susko et al. (2026) propose a solution for fair model comparison between partition models and mixture models: partition models should be evaluated using AIC scores based on marginal probabilities (mAIC), rather than those based on conditional probabilities (cAIC). In so doing, both mixture models (which naturally use marginal probabilities) and partition models (for which the cAIC is the natural AIC score, but for which an mAIC can be calculated following the procedure outlined in Susko et al. (2026)) can be compared fairly. When applied to the same simulation data used by Crotty and Holland (2022), the mAIC was shown to favour mixture models when partitions are misspecified. The mAIC allows, for the first time, mixture models to be compared directly to partition models using information theory. This opens up the possibility to answer the question of whether one type of model tends to be a better fit to empirical amino acid alignments than the other.

In this study, we sought to ask whether partition models or mixture models performed better using a collection of nine empirical datasets. For each dataset, we used best-practice methods to estimate the optimal partition models and a best-fit mixture model. We then used the newly published mAIC (Susko et al. 2026) and two additional approaches, posterior predictive simulation (Bollback 2002; Foster 2004; Brown 2014; Duchêne et al. 2018; Giacomelli et al. 2025) and a test of the robustness of the resulting phylogenetic tree inference, to compare the fit of the optimal partition and mixture models. Under the mAIC, the best mixture model substantially outperforms the best partition model in all nine of the empirical datasets we examined. We discuss the implications of this for phylogenetic inference, and the future development of models of molecular evolution in phylogenetics.

## Materials and Methods

### Empirical Datasets

We selected nine amino acid alignments from published phylogenetic studies (Table 1). These datasets were chosen to represent five major lineages: animals, plants, fungi, bacteria and archaea. They include at least hundreds of loci and 50 taxa and represent taxonomic depths from ∼59 million years (legume, Koenen et al. 2021) to ∼3.5 billion years (archaea, Moody et al. 2024).

**Table 1.** Summary of the nine datasets used in this research. The columns “Sites”, “Loci”, and “Taxa” refer to the sizes of the original datasets. “Sampled Sites in This Paper” indicates the number of alignment sites in the subsampled data used for our analyses. (*: This dataset is partitioned by protein domains)

| Clade | Sites | Loci | Taxa | Sampled Sites<br>in This Paper | Reference |
| --- | --- | --- | --- | --- | --- |
| Insects | 595033 | 2868* | 144 | 76398 | Misof et al. 2014 |
| Butterflies<br>and Moths | 749791 | 2098 | 203 | 146199 | Kawahara et al. 2019 |
| Birds | 9487705 | 14972 | 363 | 268838 | Stiller et al. 2024 |
| Green Plants | 155575 | 410 | 1187 | 152928 | One Thousand Plant<br>Transcriptomes Initiative<br>2019 |
| Legume | 325134 | 1103 | 76 | 116731 | Koenen et al. 2020 |
| Ascomycota | 562376 | 815 | 1113 | 270328 | Shen et al. 2020 |
| Budding Yeasts | 1162805 | 2408 | 343 | 189743 | Shen et al. 2018 |
| Cyanobacteria | 492937 | 1648 | 55 | 120784 | Pardo-De la Hoz et al.<br>2023 |
| Archaea | 17191 | 84 | 364 | 17191 | Dombrowski et al. 2020 |

Because some of the selected datasets are large and the estimation of partition and mixture models is computationally expensive, we randomly subsampled each dataset to 400 loci (except the green plant and archaea datasets, for which we kept all 410 and 84 loci respectively) and selected 20, 10 or 5 taxa that maximise the phylogenetic diversity (PD, Faith 1992) to investigate how the model comparison trend changes with different numbers of taxa. All subsequent analyses were then conducted on these subsampled datasets.

### Phylogenetic Model Estimation and mAIC Calculation

We conducted all phylogenetic analyses using IQ-TREE version 2.4.0 (Minh et al. 2020). Model estimation proceeded in the following steps.

First, we inferred the best partitioning schemes and best-fit models of evolution for each partition in each dataset. To do this, we used the PartitionFinder algorithm implemented in IQ-TREE, under all three alternative branch-length linkage settings: edge-equal, edge-proportional, and edge-unlinked. The optimal partition model for each of these three settings was selected based on the cAIC, and the maximum-likelihood tree was then re-optimised under this partition model.

Second, we estimated the best profile mixture model for each dataset. In principle, there are many variants of profile mixture models available in IQ-TREE, with different numbers of amino acid profiles (i.e. C10, C20, C30, C40, C50, and C60). In this study we used only the C60 model, because it is the most complex model, and because recent studies have shown that complex mixture models are generally robust to over-parameterisation in phylogenomics, because the weights of unused profiles tend to zero and have little effect on inference (Baños et al. 2024). To estimate a C60 model, we first used ModelFinder (Kalyaanamoorthy et al. 2017) to identify the best-fitting exchangeability matrix and rate heterogeneity across sites (RHAS) model (Yang 1994; Yang 1995; Soubrier et al. 2012) for each dataset, using the AIC score for model selection. Because mixture models do not partition the data, their marginal and conditional AIC scores are identical. We therefore refer to the information criterion score of a mixture model simply as its AIC throughout. We then estimated mixture models by combining the exchangeability matrix and RHAS model determined in the previous step with the C60 model implemented in IQ-TREE. During these analyses, the tree topology was fixed to that inferred under the best partition model in step 1. We estimated three variants of the C60 model, which differ in how the profiles and their weights are treated. In the first, both the 60 profiles and their weights are fixed at the published values (C60-wfix). In the second, the 60 profiles are fixed but their weights are optimised on the alignment (C60-wopt). In the third, a 61^st^ profile is added to the mixture, consisting of amino acid frequencies estimated directly from the input alignment, and the weights of all 61 profiles are optimised (C60+F, following the notation used in IQ-TREE).

All comparisons were performed on the same fixed tree topology inferred under the best partition model. Because this topology is optimised specifically for partitioned analyses, this strategy is conservative insofar as it favours partition models in subsequent model comparisons.

This procedure resulted in six scores, an mAIC score for each of the three best-fit partition models, and an AIC score for each of the three best-fit profile mixture models. These scores can be compared directly, with models that are better fit to the data receiving lower scores (Susko et al. 2026). All IQ-TREE command lines used in this study are provided (see Data and Code Availability).

### Parametric Bootstrap Measures

In addition to using the mAIC, we also sought to compare the models using a parametric bootstrap measure to assess model fit. This method was introduced by Giacomelli et al. (2025) and is a maximum likelihood equivalent of Bayesian posterior resampling (Bollback 2002). This approach asks whether data simulated under a fitted model can plausibly reproduce statistical properties of the original empirical dataset.

It should be noted here that the meaning of “parametric bootstrapping” differs from the convention in statistical testing. Parametric bootstrapping in testing is most frequently used to obtain a distribution of a test statistic under a null hypothesis. Repeated samples are generated from a model satisfying the null hypothesis and the empirical distribution of test statistics computed from these samples is treated as approximating the distribution of a test statistic of interest under the null hypothesis. In Giacomelli et al. (2025) and here, by contrast, we repeatedly generate data from a model that need not satisfy a null hypothesis; we do this separately under each model of interest and ask how closely statistics computed from the simulated data resemble those computed from the original alignment.

Giacomelli et al. (2025) used the parametric bootstrap test to assess the model adequacy of the CAT-posterior mean site frequencies (CAT-PMSF) model (Wang et al. 2018). In their framework, 100 replicate datasets were simulated under each candidate model, and the mean across-site amino acid diversity (*div*), defined as the average number of distinct amino acids observed per site, was used as the test statistic. The test simply asks whether the mean *div* value from the empirical dataset is a plausible draw of the *div* values from the simulated datasets. If it is, then the model is said to be adequate for this particular test statistic. However, if the empirical mean *div* value falls outside the distribution of mean *div* values from the simulated data, then it can be concluded that the model inadequately captures the process which generated the data.

The approach of Giacomelli et al. (2025) is attractive, but using the mean of the *div* statistic has two important limitations. First, it considers only the presence or absence of amino acids at each site and ignores their relative frequencies. Second, as a single summary value, the mean only crudely captures differences in the full distribution of values across sites in the dataset. To address these limitations, we extend the approach of Giacomelli et al. (2025) in two directions. First, we replace the diversity of each site (their *div*) with the Shannon entropy of each site (Shannon 1948). The Shannon entropy is attractive because it accounts for variation in the frequencies of amino acids at each site, not just their presence or absence. Second, instead of comparing the mean of the distribution as the test statistic, we employ the two-sample Cramér–von Mises test (CvM test, Anderson 1962) to directly compare the full distributions of the *div* or entropy values at each site in the simulated datasets to those in the original empirical datasets. In this version of the parametric bootstrap, we obtain a distribution of CvM statistics (W^2^ values) - one per simulated replicate - where each value quantifies how closely that replicate’s per-site entropy distribution resembles the empirical one. A W^2^ value of zero would indicate a perfect match, so a model that adequately captures the generative process should produce a distribution of W^2^ values concentrated near zero. A model that fails to capture the true process will yield systematically larger W^2^ values. Models can therefore be compared by examining these distributions: the preferred model is the one whose simulated datasets are, on average, most similar to the empirical data, and so whose CvM statistic distribution is closer to zero.

As mentioned above, our parametric bootstrap does not generate data under a null hypothesis, so no threshold for an acceptable W² can be defined. Furthermore, because W² measures the distance between a simulated replicate and the empirical alignment, its value for the empirical data is zero by construction, and so it provides no reference against which the simulated distributions could be judged. The CvM test therefore ranks the candidate models relative to one another but cannot establish that any of them is adequate, and it will identify a preferred model even when all of the candidates fit the data poorly. In this sense, the intended use of the CvM test here is similar to the use of the AIC in phylogenetic model selection - both measures provide a rank-ordering of models, but neither gives information on absolute model adequacy (Shepherd and Klaere 2019). The two approaches are complementary: the mean from the data can be compared with the distribution of means simulated from a model to test whether the model-based mean can be rejected for the data. The CvM statistic provides a finer relative comparison, but in order to do this it uses a statistic for each simulated data set that depends on both the simulated data and the empirical data, not just the empirical data.

To allow for direct comparison with the method of Giacomelli et al. (2025), we implemented four parametric bootstrap measure strategies in this study: mean *div*, mean entropy, CvM test on *div* distributions, and CvM test on Shannon entropy distributions. For each estimated partition or mixture model, we simulated 100 replicate datasets using AliSim (Ly-Trong et al. 2022). For each strategy, we computed the corresponding statistic for the simulated datasets and compared them with those derived from the empirical data to assess model adequacy.

### Model Robustness Test

Tree inference is a central goal of phylogenetic research. To that end, we implemented a third way to compare partition and mixture models, which uses the robustness of tree inference as the test statistic. To evaluate tree robustness, we inferred the tree topologies using a leave-one-taxon-out jackknife approach. For each 20-taxon alignment, we performed jackknife resampling by sequentially removing one sequence at a time, generating twenty 19-taxon sub-alignments. Because this test is computationally intensive, we performed it under the best-fitting mixture and partition models as identified by the information criteria (Fig. 1). Among the C60 variants, C60+F was consistently the best across all nine datasets. Among the partition models, the mAIC favoured the edge-proportional model whereas the cAIC favoured the edge-unlinked model. For each sub-alignment, we therefore inferred trees under the C60+F model and under the edge-proportional and edge-unlinked partition models. Each resulting 19-taxon tree was then compared to the corresponding pruned version of the tree inferred from the full 20-taxon alignment. A model is considered more robust the more similar the 19−taxon trees are to the original 20-taxon tree. In other words, we prefer models that estimate robust tree topologies when the dataset is perturbed in a relatively minor way (in this case, the removal of one taxon).

**Fig. 1.**
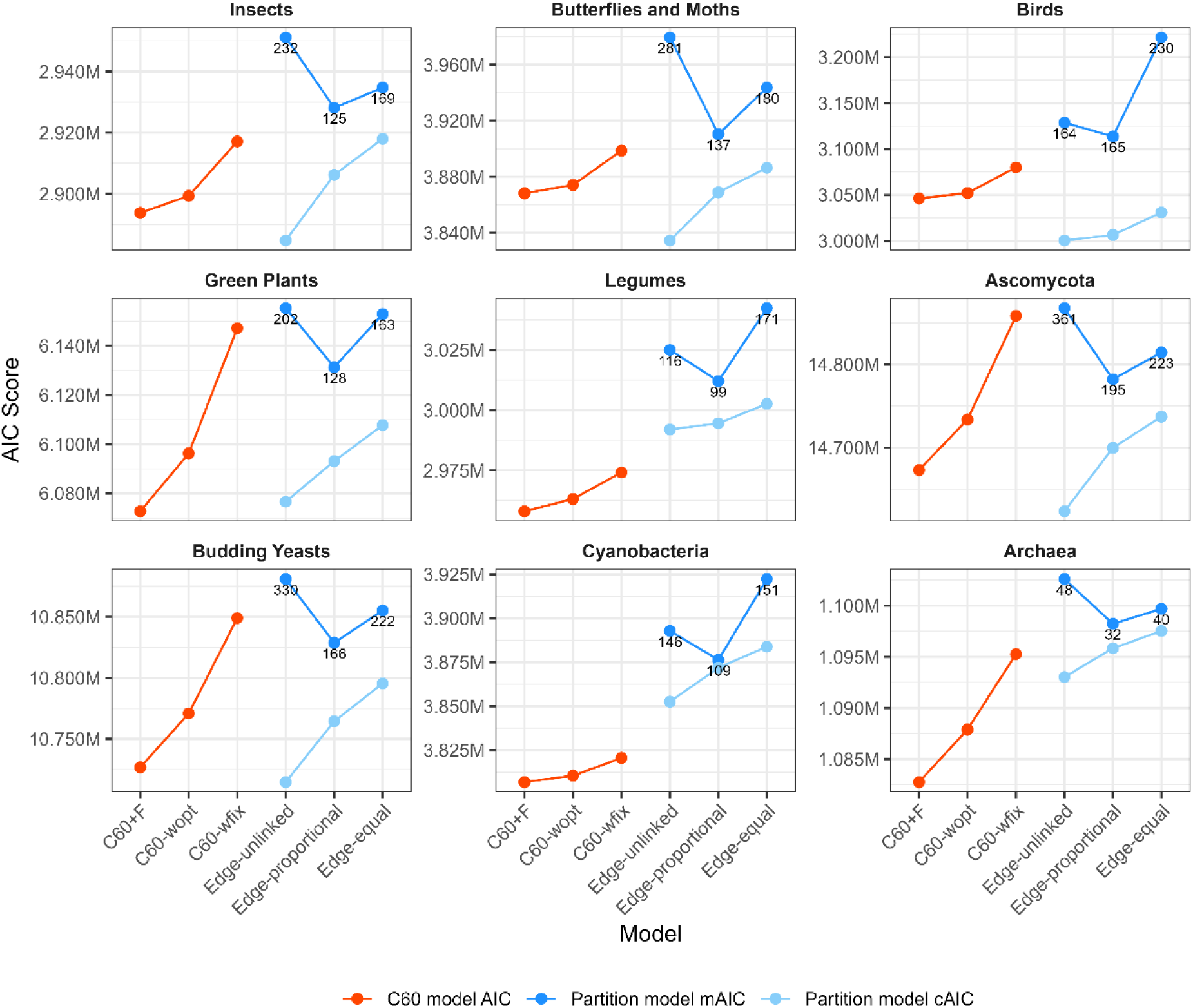
AIC scores of C60 and partition models on 20-taxon datasets. The x-axis shows the six model types, and the y-axis shows the corresponding AIC values. Numbers above the partition models indicate the optimal number of partitions inferred by PartitionFinder. Note that the AIC for the C60 model (red) and the mAIC for the partition model (dark blue) are directly comparable. Neither can be compared directly to the cAIC for the partition model (light blue). The introduction explains why this is the case. The cAIC is shown purely to demonstrate that the cAIC and mAIC make different predictions about which of the three partition models best fits each dataset.

Tree topological differences were quantified using the Lin-Rajan-Moret (LRM) distance (Lin et al. 2012), rather than the widely used Robinson-Foulds (RF) distance (Robinson and Foulds 1981). The RF distance simply counts the number of discordant splits. A critical limitation is that a small structural change, such as moving a single terminal branch to a different position, can result in a large RF distance. The LRM distance addresses this issue. The LRM and RF distances between two trees can both be viewed as constructing a one-to-one mapping of splits in the first tree to splits in the second, such that the sum of split distances is minimal. For RF the distance between splits is simply 0 or 1 according to whether they are the same or not. The LRM split distance is more sensitive: it is the minimum number of taxa that need to be removed in order to make the two splits the same. Under this scheme, moving one taxon to a different place in the tree will lead to a relatively small addition to the LRM distance. All tree pruning and LRM distance calculations were performed in Cogent3 (Knight et al. 2007).

### Summary of Model Comparison

Overall, we compared partition and mixture models using three complementary criteria. For each dataset, a model was considered to perform better if it yielded a lower mAIC score, showed a closer fit in the parametric bootstrap analyses, and produced tree topologies that were more robust to taxon removal.

## Results

### mAIC strongly favours C60 profile mixture models over best-fit partition models

Figure 1 presents the AIC scores of the C60 models and the mAIC and cAIC scores of the partition models fitted to our nine empirical datasets subsampled to 20 taxa. Note that the only valid comparison in Figure 1 is between the AIC scores for the C60 models and the mAIC scores for the partition models (i.e. the red dots and dark blue dots can all be compared to one another). The cAIC scores for partition models cannot be compared directly to the other AIC scores (i.e. the light blue dots can be compared to each other, but not to the red or dark blue dots). Figure 1 shows that the C60 models consistently outperform the partition models across all datasets. Among C60 model variants, the AIC scores follow a consistent ranking: C60+F < C60-wopt < C60-wfix. For partition models, the edge-proportional model always has the best mAIC score, and also always has the smallest number of partitions. In the green plant, Ascomycota and budding yeast datasets, edge-proportional partition models outperform the C60-wfix models, but both are substantially outperformed by the other C60 models. Notably, cAIC scores of partition models consistently rank edge-unlinked < edge-proportional < edge-equal, are always lower than their corresponding mAIC scores, and are sometimes lower than the AIC scores of the C60 models. This serves as a useful reminder that comparing the cAIC of a partition model to the AIC of a mixture model can be very misleading (Crotty and Holland 2022; Susko et al. 2026).

The results for the 10- and 5-taxon subsampled datasets are qualitatively identical to the 20-taxon subsets. They also show that the mAIC favours the best C60 model (Supplementary Fig. S1 and S2) over the best partition model in every dataset. However, the difference in AIC scores between the partition models and C60 models is smaller. In the smallest dataset (archaea, with 84 loci and 5 taxa), only the C60+F model yields a lower AIC score than the mAIC score of the edge-proportional and edge-unlinked partition models.

### C60 models outperform partition models in parametric bootstrap measures

Figure 2 shows the distributions of W² values, the two-sample Cramér–von Mises (CvM) statistic, comparing the distribution of site-wise Shannon entropy values for each empirical dataset against 100 simulated datasets (see Materials and Methods). Lower W² values indicate that the per-site Shannon entropy of simulated datasets more closely matches that of the original empirical data. Across seven of the nine datasets (except the budding yeast and Ascomycota datasets), C60 models fit the data better than the partition models, with C60-wfix models performing the best. In contrast, for the budding yeast and Ascomycota datasets, the edge-unlinked partition models provide the best fit, followed by the C60-wopt and the C60+F models while the C60-wfix models perform poorly. Among the partition models, edge-unlinked models outperform others across all nine datasets. Results based on mean Shannon entropy values (Fig. 3) supplement these findings: the C60-wfix models produce the smallest Shannon entropy distributions, which are often the closest distributions to the mean Shannon entropy of the empirical alignment (shown with a dashed vertical line). Other models tend to overestimate the empirical entropy. However, in the Ascomycota and budding yeast datasets, the C60-wfix model substantially underestimates entropy, and the edge-unlinked partition model overestimates the Shannon entropy but to a lesser extent. These observations are concordant with the results from the CvM distributions shown in Figure 2.

**Fig. 2.**
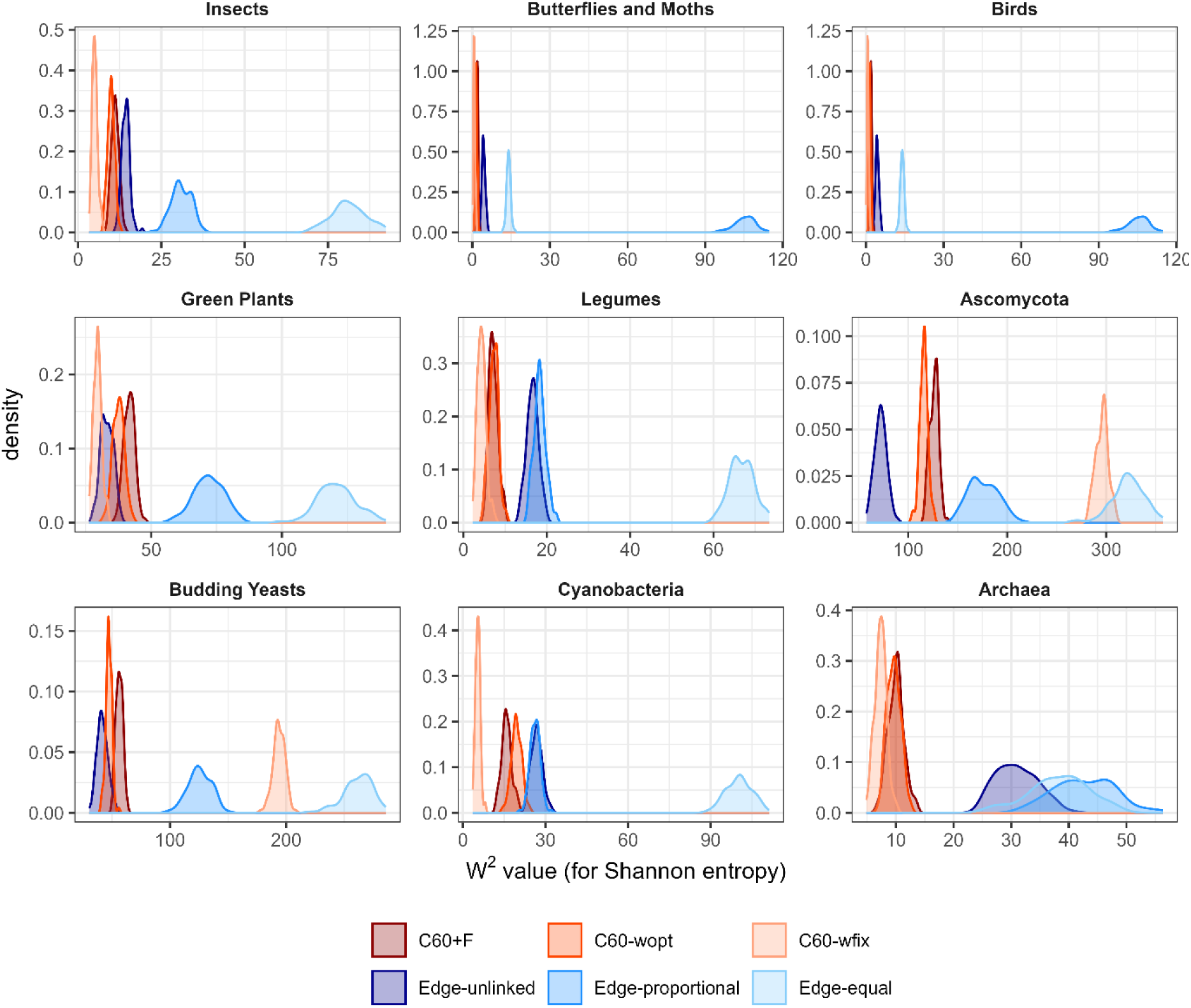
Distributions of CvM test statistics (W² values) for Shannon entropy of simulated datasets under different C60 and partition models.

**Fig. 3.**
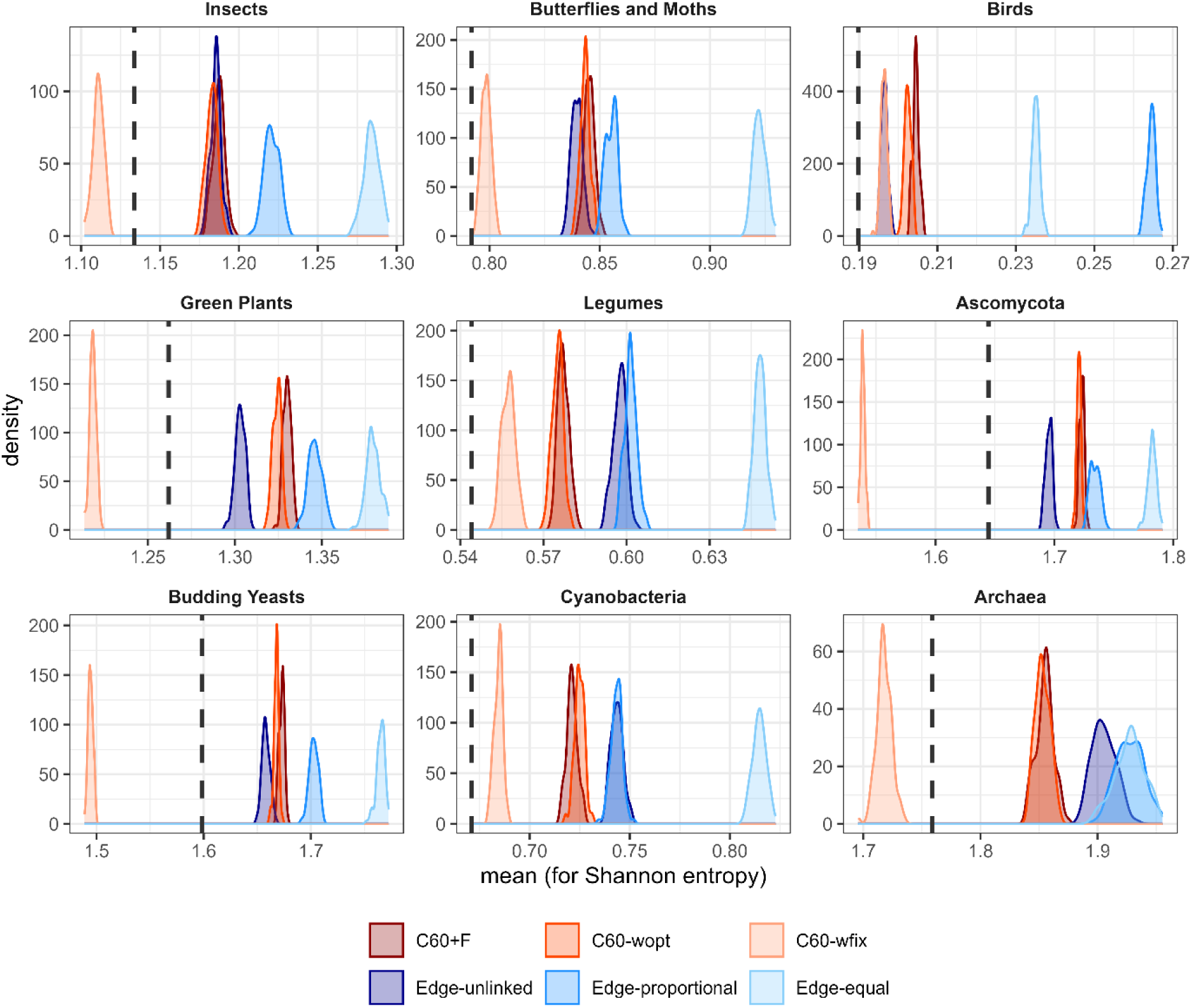
Distributions of mean Shannon entropy of simulated datasets under different C60 and partition models. The dashed line indicates the mean Shannon entropy of sites in the real data.

The results for CvM and mean-based parametric bootstrap simulations using *div* values agree with those from the Shannon entropy analyses shown in Figures 2 and 3 (see Supplementary Fig. S3 and S4). Across all datasets, the *div* values of the C60-wfix and C60-wopt models are the closest to the empirical data. Similarly, for the Ascomycota and budding yeasts datasets, the C60-wfix model underestimates *div*, whereas the C60-wopt model outperforms the partition model. Among the partition models, edge-unlinked models again perform the best, consistent with the Shannon entropy results.

Additionally, supplementary Figures S5–S13 present cumulative distribution functions (CDFs) of site-wise Shannon entropy for nine datasets. For each model, the site-wise Shannon entropy values from all 100 simulated replicates were pooled into a single distribution (i.e., for an alignment with N sites, this yields 100 × N entropy values), and the resulting CDF was compared against the CDF of the N site-wise entropy values from the empirical alignment. These CDF plots corroborate the findings from Figures 2 and 3. Among the models, C60-wfix produces entropy distributions closest to the empirical CDF in most datasets, except the budding yeast and Ascomycota datasets, while the C60+F, C60-wopt and partition models tend to overestimate entropy at higher values. For the Ascomycota and budding yeast datasets, the edge-unlinked model provides a closer fit to the empirical CDF, as the C60-wfix underestimates entropy. These results are consistent with the CvM results reported above.

### C60 and partition models are both similarly robust

The distribution of LRM scores comparing the 20-taxon alignment tree to the corresponding leave-one-taxon-out jackknife subalignment trees (Fig. 4) shows that different models are favoured for different datasets. For most datasets, the best mixture model (C60+F) performs very similarly to the best partition model (the edge-proportional model). For the butterflies and moths and legume datasets, the C60+F model performs better than the best partition model. But for the budding yeasts dataset, the edge-proportional partition model performs better than the C60+F model.

**Fig. 4.**
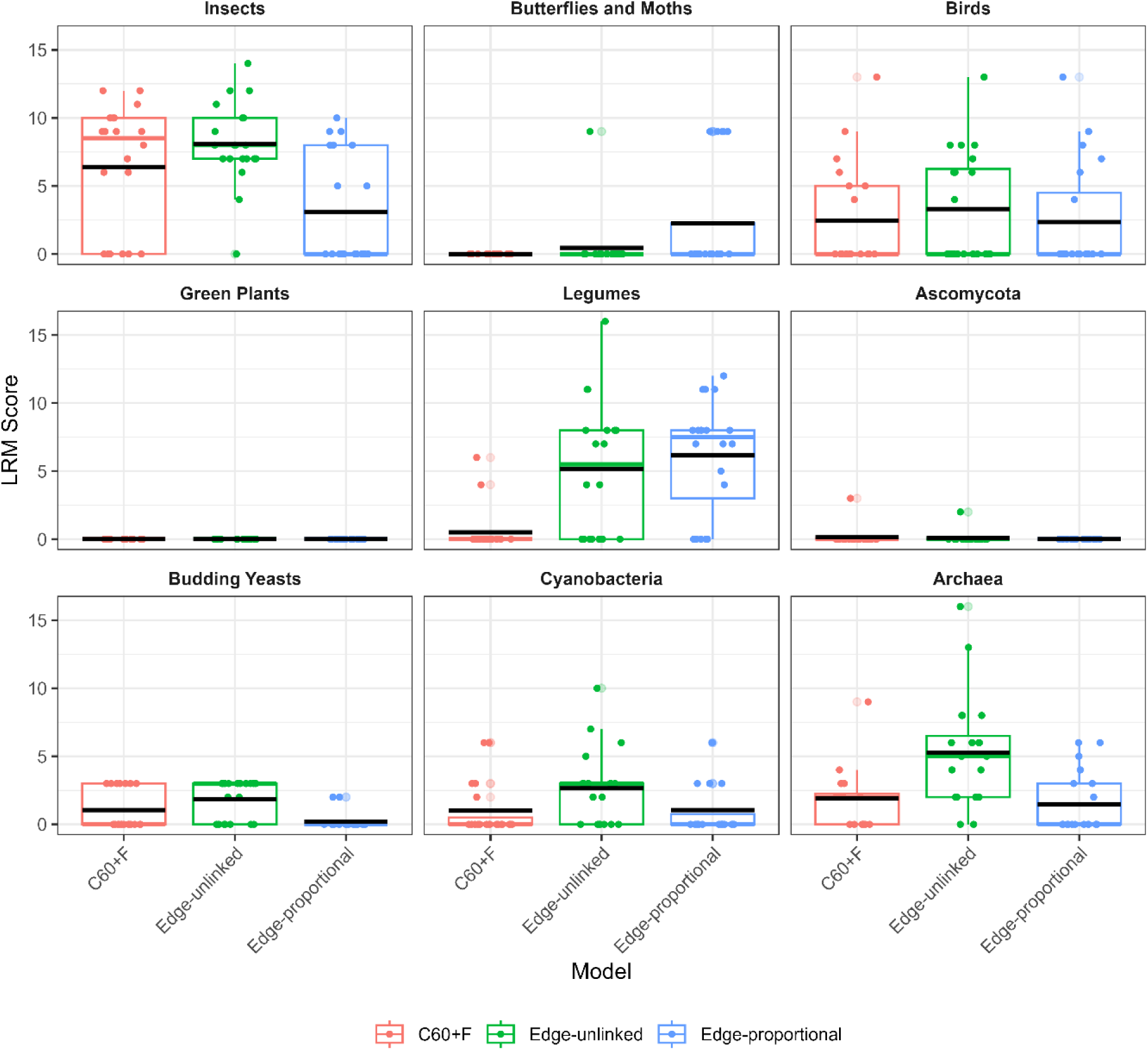
The distribution of LRM scores comparing the 20-taxon alignment tree to the corresponding leave-one-taxon-out jackknife subalignment trees under different models. Lower LRM scores indicate greater similarity to the reference tree, and therefore greater robustness. The black horizontal line denotes the mean of the LRM scores of each model.

## Discussion

In this study, we evaluated partition models and the C60 profile mixture models, a widely used class of mixture models, under various settings. We fitted these models to nine empirical datasets representing diverse taxonomic groups and compared their fits using three complementary approaches. Across all datasets, C60 models consistently outperformed partition models by mAIC, and in most datasets by parametric bootstrap as well, while the robustness test gave a more equivocal result.

Across all comparisons, the best-performing C60 models consistently achieved lower mAIC scores than the best partition models. According to Burnham and Anderson (2004), a difference of about 10 AIC units is generally considered strong evidence in favor of the model with the lower score. Although the exact threshold may differ for mAIC, the differences observed here are far larger: the best C60 models improve upon the best partition models by thousands of mAIC units, or even more. Notably, the advantage of the C60 models became more pronounced as dataset size increased (Figs. 1, S1, and S2). For the smallest dataset considered here, the 5-taxon archaea dataset, the best partition model slightly outperformed the weakest C60 variant. However, this pattern was not observed in datasets with larger numbers of taxa or loci. Given that empirical phylogenomic datasets typically include far more than five taxa, these results suggest that C60 mixture models provided a consistently better fit to the empirical data than the best partition model.

Besides mAIC, which directly assesses all aspects of model fit by using the likelihood of the observed data, we employed two additional approaches, parametric bootstrap measures and robustness tests, to further study model fit. The parametric bootstrap has been widely applied in phylogenetic model selection (Emerson et al. 2001; Quang et al. 2008; Giacomelli et al. 2025). We extended the parametric bootstrap framework of Giacomelli et al. (2025) and found that the C60 mixture models better reproduce the empirical distributions of site-wise amino acid diversity and Shannon entropy than the partition models for most datasets. The partition models tend to overestimate the amino acid diversity per site, because all sites within each data block share the same substitution profile. C60 models are specifically designed to relax this constraint, modelling the tendency of many sites in a protein to accept only a small subset of the 20 available amino acids. Partition models, on the other hand, are forced to estimate a single average amino acid profile for each partition, resulting in appreciable frequency for most amino acids and, consequently, larger predicted entropy and diversity in each site. Notably, in a few cases, the C60-wfix model underestimated the amino acid diversity, while other tested models never did, likely because the predefined profiles in the C60-wfix do not reflect the composition patterns of many of the empirical datasets we analyse.

In the model robustness test, C60+F mixture models and edge-proportional partition models both exhibited similar robustness across datasets. Our results indicate that no single modelling strategy was uniformly more or less robust than another. The advantage of the C60 model over the partition models is therefore less pronounced here than it is under the model fit criteria, but the test provides a complementary dimension of model evaluation, one based on tree inference rather than on model fit statistics alone. It is worth being explicit that the robustness test measures how consistently a model recovers the same topology when the data are perturbed, not whether the topology it recovers is correct. Two models may be equally robust while recovering different trees, with one consistently recovering the correct tree and the other consistently recovering the incorrect tree, much as the bootstrap support can be high for incorrect relationships when the model is misspecified (Huang et al. 2021). The similar robustness of mixture and partition models should accordingly not be taken to indicate that the two classes of model are equally suitable for phylogenetic inference. So, while robustness provides useful information on model performance, one might still choose a less robust model if other statistics indicate that that model provides a more adequate description of the data in hand.

Over the past decades, comparisons between partition and mixture models have relied on downstream analyses, such as cross-validation (Susko and Roger 2020) and bootstrap support (Jhwueng et al. 2014; Whelan and Halanych 2017) and posterior predictive simulation (Giacomelli et al. 2025; Pisani et al. 2026). While these approaches have provided valuable insights into model behaviour, their high computational costs and complex workflows limit their routine application in empirical phylogenetic studies. In practical phylogenetic analyses, there remains a lack of efficient and easily applicable frameworks for comparing partition and mixture models. In this study, we addressed this gap by employing the newly introduced information criterion mAIC, which enables direct and computationally tractable comparisons between partition and mixture models.

For comparisons among different branch length settings within partition models, edge-proportional models have been suggested to provide the best fit (Duchêne et al. 2020). Duchêne et al. (2020) report that BIC correctly supports edge-proportional models, due to its stronger penalty for additional parameters, whereas AICc tended to support edge-unlinked models. Our results (Supplementary Figs. S14-S16) are consistent with these previous findings. Among information criteria based on conditional likelihoods, BIC favoured edge-proportional models but cAIC and AICc favoured edge-unlinked models. Interestingly, mAIC favours edge-proportional models rather than edge-unlinked models, in agreement with BIC. This pattern suggests that mAIC may offer advantages over cAIC and AICc in selecting partitioning schemes. Meanwhile, the tree robustness test (Fig. 4) also supports edge-proportional models rather than edge-unlinked models.

According to our results (Fig. 1; Supplementary Figs. S1 and S2), mAIC scores are strongly correlated with the number of partitions identified by PartitionFinder. This is likely because exchangeability matrices and rates are estimated on partition-specific data blocks, which makes them less generalisable and leads to poorer marginal likelihoods when evaluated across the full alignment. To reconcile the difference in model preferences between mAIC and cAIC, we adopt the reasoning proposed by Susko et al. (2026) who point out that the cAIC tends to favour the partition models when partitions are misspecified, even with infinite data. If the marginal model is correctly specified, the global parameters, including tree topology, will also be correct. Thus, model selection procedures should be able to recognise correct marginal specifications. This implies that if the research focuses on partition-specific properties like exchangeability matrices or rates, the cAIC is the more appropriate metric with which to select a model. But if the research is interested in global parameters like the shared tree topology, the mAIC should be used.

It is noteworthy that our mAIC and parametric bootstrap results lead to different preferences among partition models: the mAIC favours edge-proportional models, whereas the parametric bootstrap favours edge-unlinked models (Fig. 2, 3, S3-S4 and S5-S13). This discrepancy is likely attributable to two fundamental differences between the two approaches. First, they summarize model fit in different ways: the mAIC evaluates the fit of the model to the data as a whole, capturing a wide range of aspects of model fit, whereas the parametric bootstrap as implemented here focuses on a single summary of the data, the distribution of site-wise entropy values. This limitation of the parametric bootstrap is highlighted by its disagreement with the tree robustness results: the parametric bootstrap clearly prefers the edge-unlinked partition model over the edge-proportional model for most datasets, but the tree robustness test shows the opposite trend. Second, and perhaps more importantly, information criteria penalize model complexity, whereas the parametric bootstrap does not. Edge-unlinked models have substantially more parameters than edge-proportional models, giving them greater flexibility to reproduce specific features of the empirical data such as entropy distributions. This difference likely explains why the edge-unlinked models perform better than the edge-proportional models in the parametric bootstrap but not under mAIC.

These observations again highlight the importance of choosing model selection criteria that are appropriate for the question at hand. In principle, many different summary statistics could be used to compare simulated with real data, and no single choice is uniquely correct. What matters is that the statistic reflects the quantity one is actually trying to estimate. For example, if the aim of a study was to accurately reproduce site-wise entropy or diversity values, then a parametric bootstrap based on precisely these values, or the use of an information criterion more suited to measuring the fit of local parameters such as the cAIC (Susko et al. 2026), would be appropriate model selection criteria. However, if the aim of a study was to accurately reconstruct a global parameter such as the concatenated tree topology, which is often the primary interest of empirical biologists, then an information criterion more suited to measuring the fit of global parameters, such as the mAIC, would be the more appropriate model selection criterion. The tree robustness test operates on the same global parameter and provides a useful complement, but as noted above, it measures the stability of an inferred topology rather than its accuracy. Thus, where two models are indistinguishable in terms of model fit (e.g. mAIC), one should prefer the model with higher robustness.

For C60 model variants, the AIC, AICc and BIC values all follow the order C60+F < C60-wopt < C60-wfix across all datasets (Supplementary Fig. S17-S19), indicating that C60+F fits the data better. This is likely because C60+F incorporates empirical amino acid frequencies estimated from the input alignment, whereas C60-wopt only optimises the weights of pre-defined profiles, and C60-wfix uses fixed, pre-defined profile weights without any optimisation. However, a previous study suggests that when exchangeability matrices are misspecified, empirical frequency (+F) can sometimes lead to better likelihoods but incorrect tree topologies (Baños et al. 2024). Similarly, our parametric bootstrap analyses suggest that the performance of C60-wopt and C60+F models is close.

Our primary analyses focused on C60 profile mixture models, which are complex mixture models routinely applied in most phylogenomic work. Previous work has shown that profile mixture models are generally robust to over-parameterisation because unused profiles tend to receive negligible weights (Baños et al. 2024; Pisani et al. 2026). This suggests that the advantages we observe for the C60 models are likely to extend to other mixture models of equal or greater complexity, such as the UDM models (Schrempf et al. 2020) or mixture-within-partition approaches (Redmond and McLysaght 2021; Susko et al. 2026).

However, the same conclusion cannot be assumed for simpler mixture models. For instance, we repeated our mAIC comparison using the C10 models on all nine datasets (Supplementary Fig. S20). The C10+F model still achieved a lower AIC score than the mAIC score of the best partition model in every dataset, but in six of the nine datasets, the C10-wopt and/or C10-wfix models were outperformed by the best partition model. Our C10 results are consistent with those of Pisani et al. (2026), who independently found in simulations that C10-based models were the worst performing of the heterogeneous models they tested, and attributed this to the limited number of profiles available to C-series models. Additionally, the partition models compared in this study were selected using the cAIC, as implemented in PartitionFinder. We have recently implemented the mAIC within PartitionFinder and shown that the partition models it selects have better mAIC scores than those selected using the cAIC (Ren et al. 2026). Partition models estimated in this way might therefore compare more favourably than those reported here, which would narrow the margin over the simpler mixture models, such as C10, in particular. We note, however, that this is unlikely to affect our main conclusion: the best C60 models improve on the best partition models by thousands of mAIC units (Fig. 1).

A limitation of our study is that each dataset was reduced to at most 400 loci and 20 taxa before analysis. Fitting partition and mixture models to the full alignments, and in particular fitting the C60 models, was computationally prohibitive at the scale of this study, and we therefore neither repeated the random subsampling of loci nor extended the analyses to larger subsets. However, one aspect of our design does bear on the robustness of the comparison: we repeated the mAIC analyses on 20-, 10- and 5-taxon subsets, and found that the advantage of the C60 models over the partition models became more pronounced as dataset size increased. Analyses of larger subsets would therefore be expected to strengthen rather than weaken our conclusion, although a systematic assessment of how these model comparisons vary across repeated subsamples and larger datasets would be a valuable direction for future work.

This study is the first to show that mixture models often perform better than partition models when evaluated by an appropriate information criterion. Two complementary assessments, a parametric bootstrap and a test of tree robustness, are broadly consistent with this conclusion. We hope that mAIC will become a valuable tool for comparing complex models in current and future phylogenetic analyses.

## Data and Code Availability

All code used in this study, including scripts for data subsampling, IQ-TREE 2 command-line execution, parametric bootstrap measures, and model robustness analyses, is available in our GitHub repository at: https://github.com/ChangsenJiang/Partition-vs-Mixture/tree/main and in the Dryad repository at: https://doi.org/10.5061/dryad.8cz8w9h73.

The raw sequence alignments analysed in this study, the raw data underlying all figures, the IQ-TREE output files used for the analyses and the IQ-TREE program we used (version 2.4.0) are also available in these repositories.

## Supplementary Materials

Supplementary material is available at [supplementary file link]

