## Supplementary Materials for "A New Information Theoretic Approach Shows that Mixture Models Outperform Partition Models for Phylogenetic Analyses of Amino Acid Data"

### Supplementary Figures

**Fig. S1.** AIC scores of C60 and partition models on 10-taxon datasets. The x-axis shows the six model types, and the y-axis shows the corresponding AIC values. Numbers above the partition models indicate the optimal number of partitions inferred by PartitionFinder.

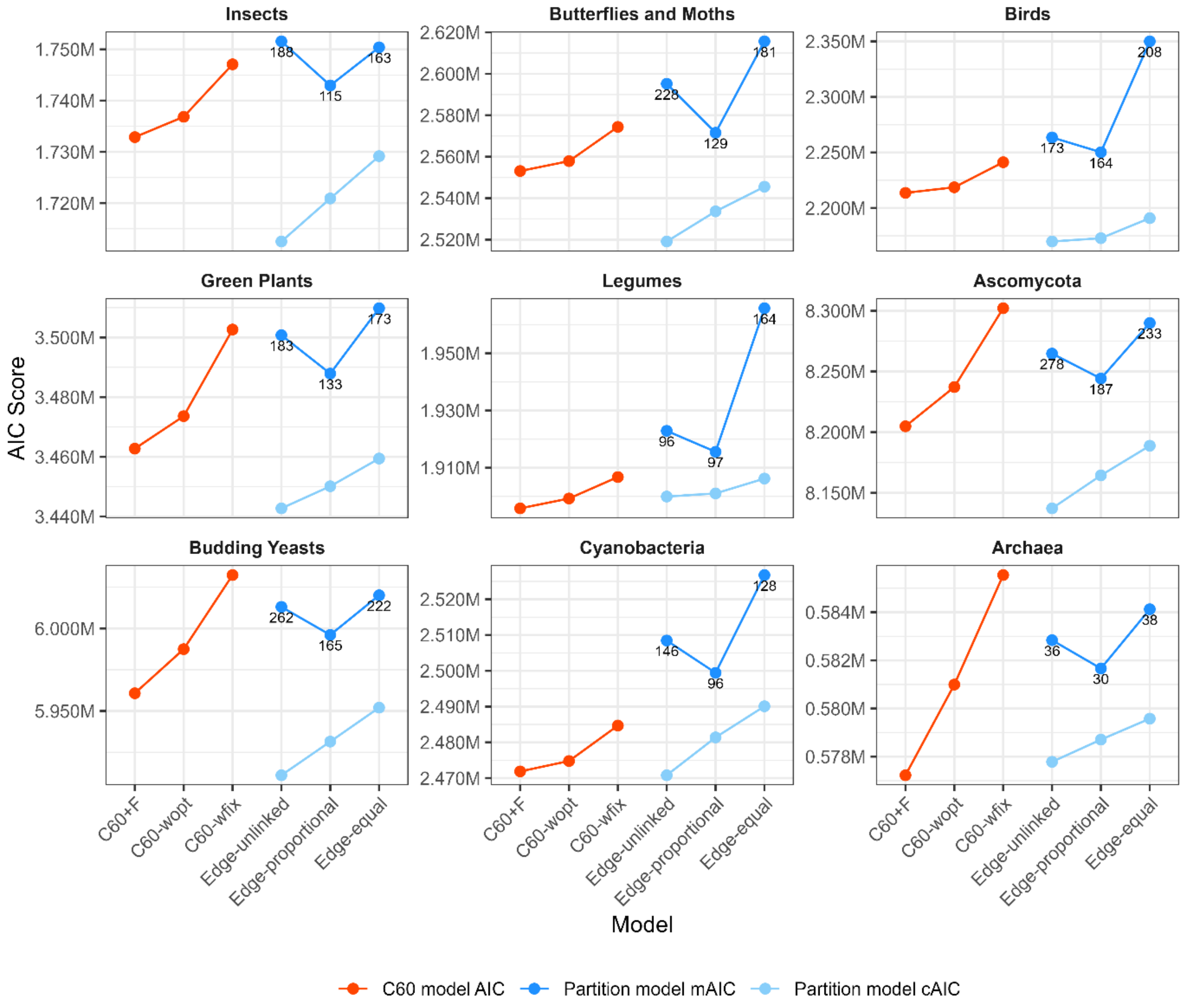

**Fig. S2.** AIC scores of C60 and partition models on 5-taxon datasets. The x-axis shows the six model types, and the y-axis shows the corresponding AIC values. Numbers above the partition models indicate the optimal number of partitions inferred by PartitionFinder.

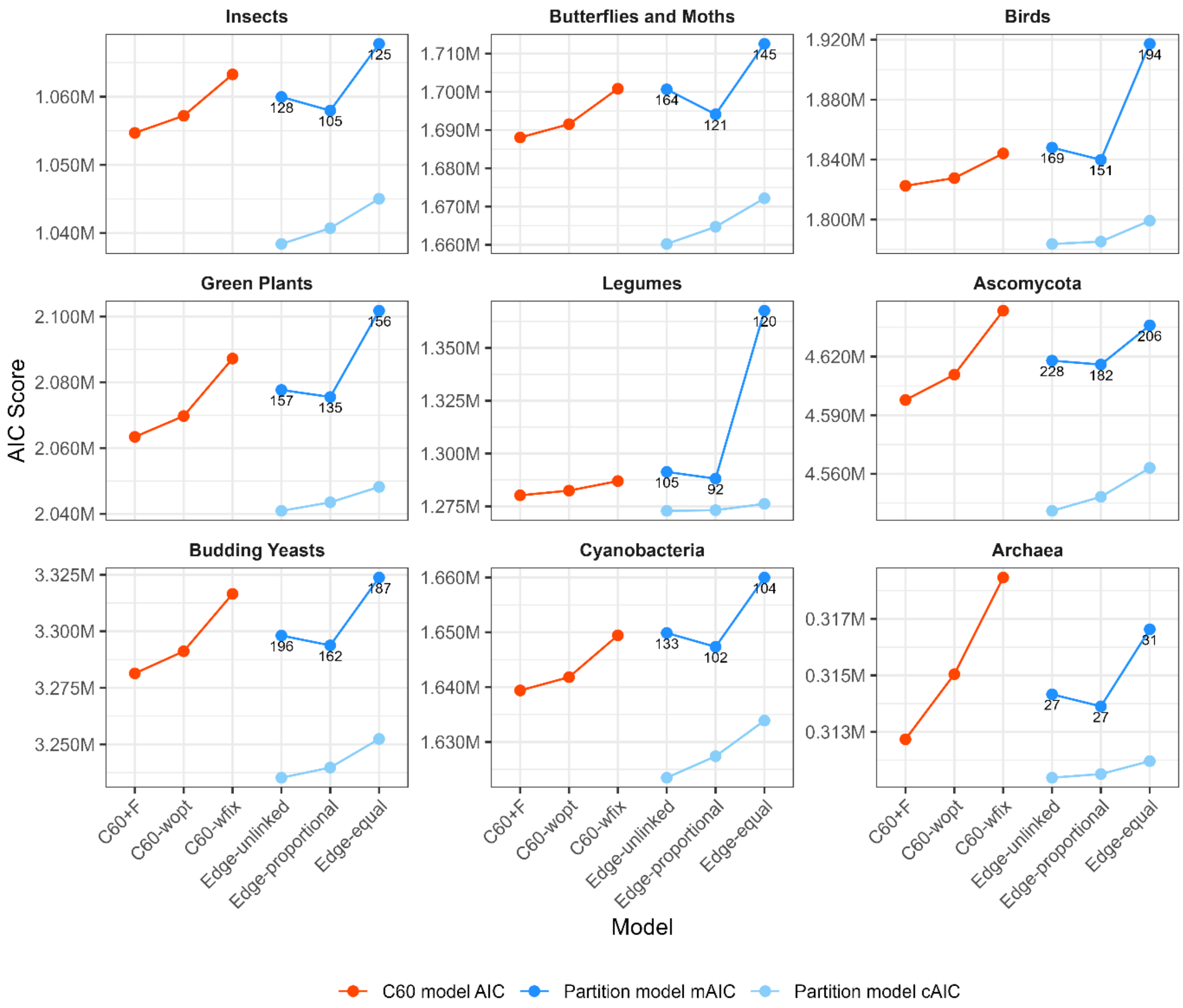

**Fig. S3.** Distributions of CvM test statistics (W² values) for *div* of simulated datasets under different C60 and partition models.

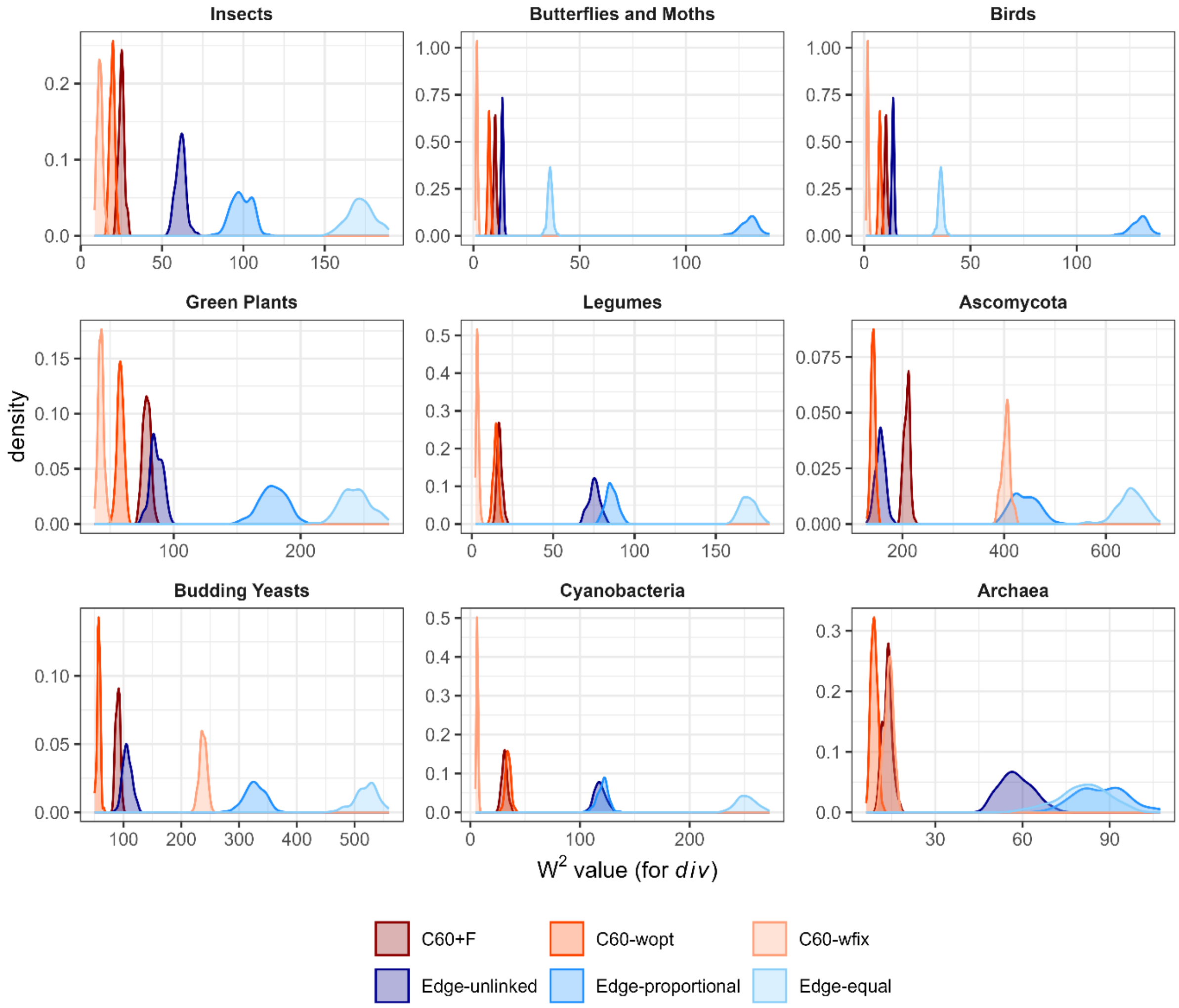

**Fig. S4.** Distributions of mean *div* of simulated datasets under different C60 and partition models. The dashed line indicates the mean *div* of sites in the real data.

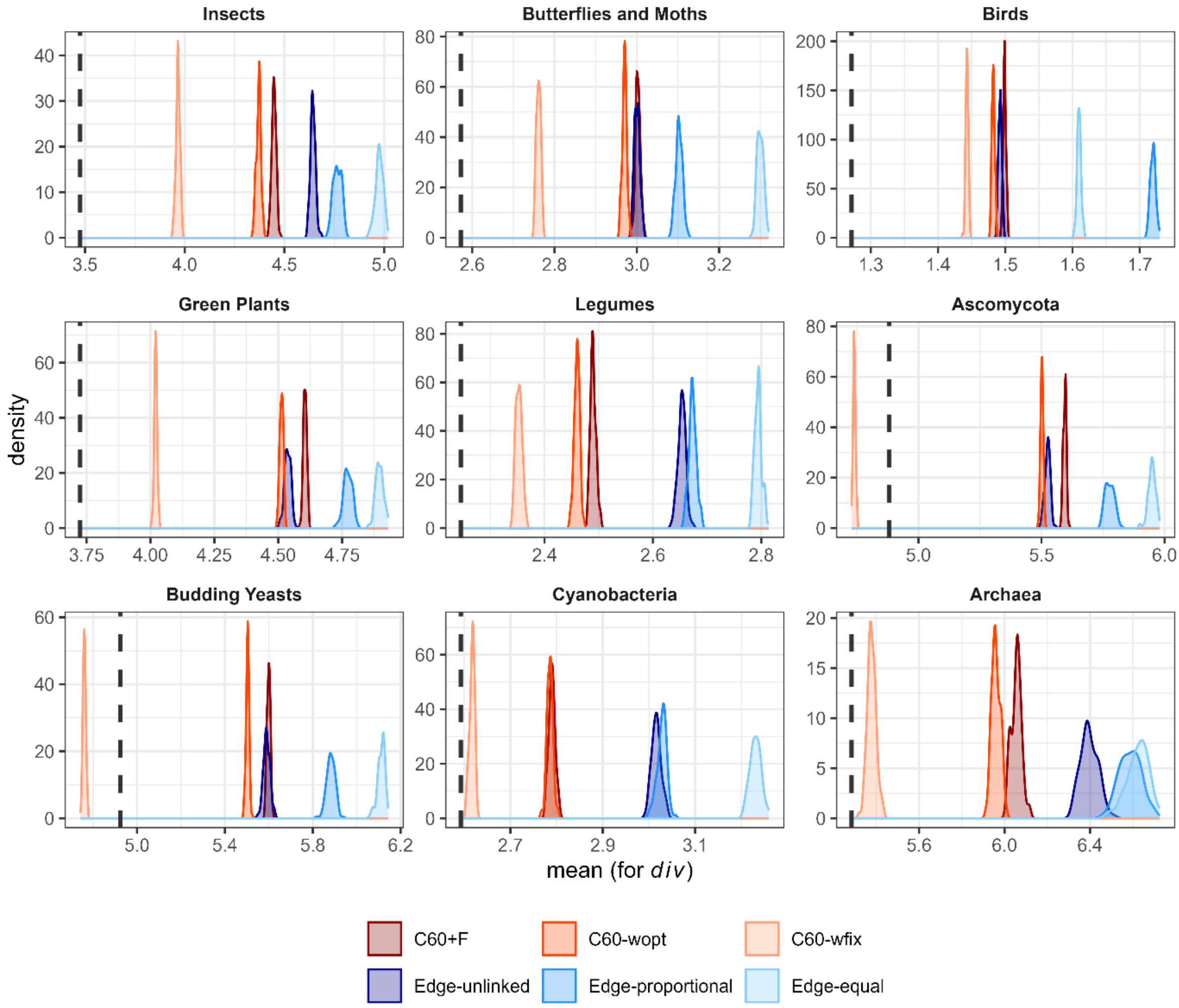

**Fig. S5.** Cumulative distribution function of site-wise Shannon entropy for 100 simulated datasets under six models compared to the raw insects dataset (subsampled to 20 taxa). For each of the six models, site-wise Shannon entropy values from 100 simulated replicate datasets were pooled into a single distribution, and the resulting CDF (coloured lines) is compared against the CDF of the empirical alignment (black line). Note that some model CDFs may closely overlap and appear visually indistinguishable in the plot.

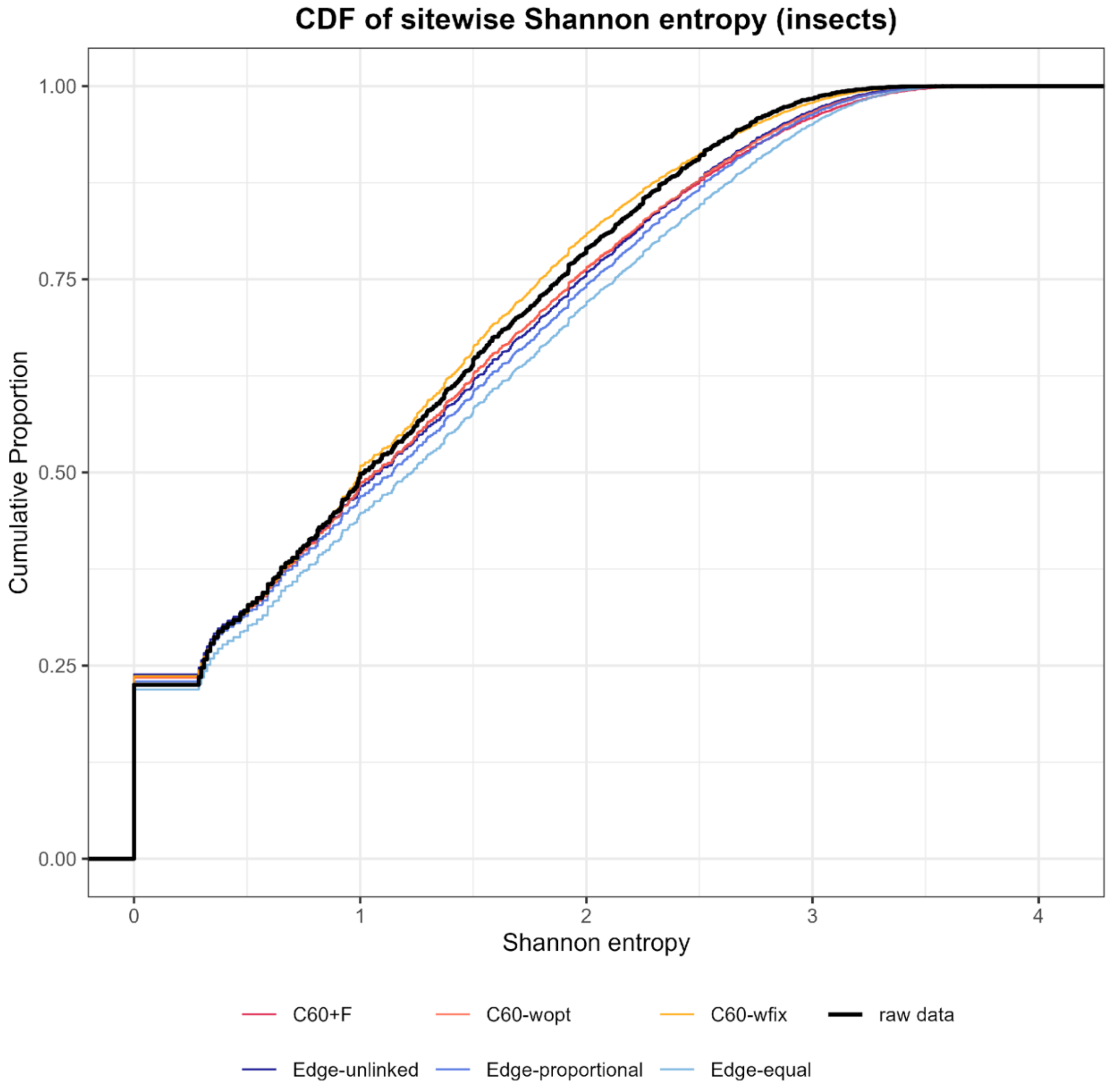

**Fig. S6.** Cumulative distribution function of site-wise Shannon entropy for 100 simulated datasets under six models compared to the raw butterflies and moths dataset (subsampled to 20 taxa). For each of the six models, site-wise Shannon entropy values from 100 simulated replicate datasets were pooled into a single distribution, and the resulting CDF (coloured lines) is compared against the CDF of the empirical alignment (black line). Note that some model CDFs may closely overlap and appear visually indistinguishable in the plot.

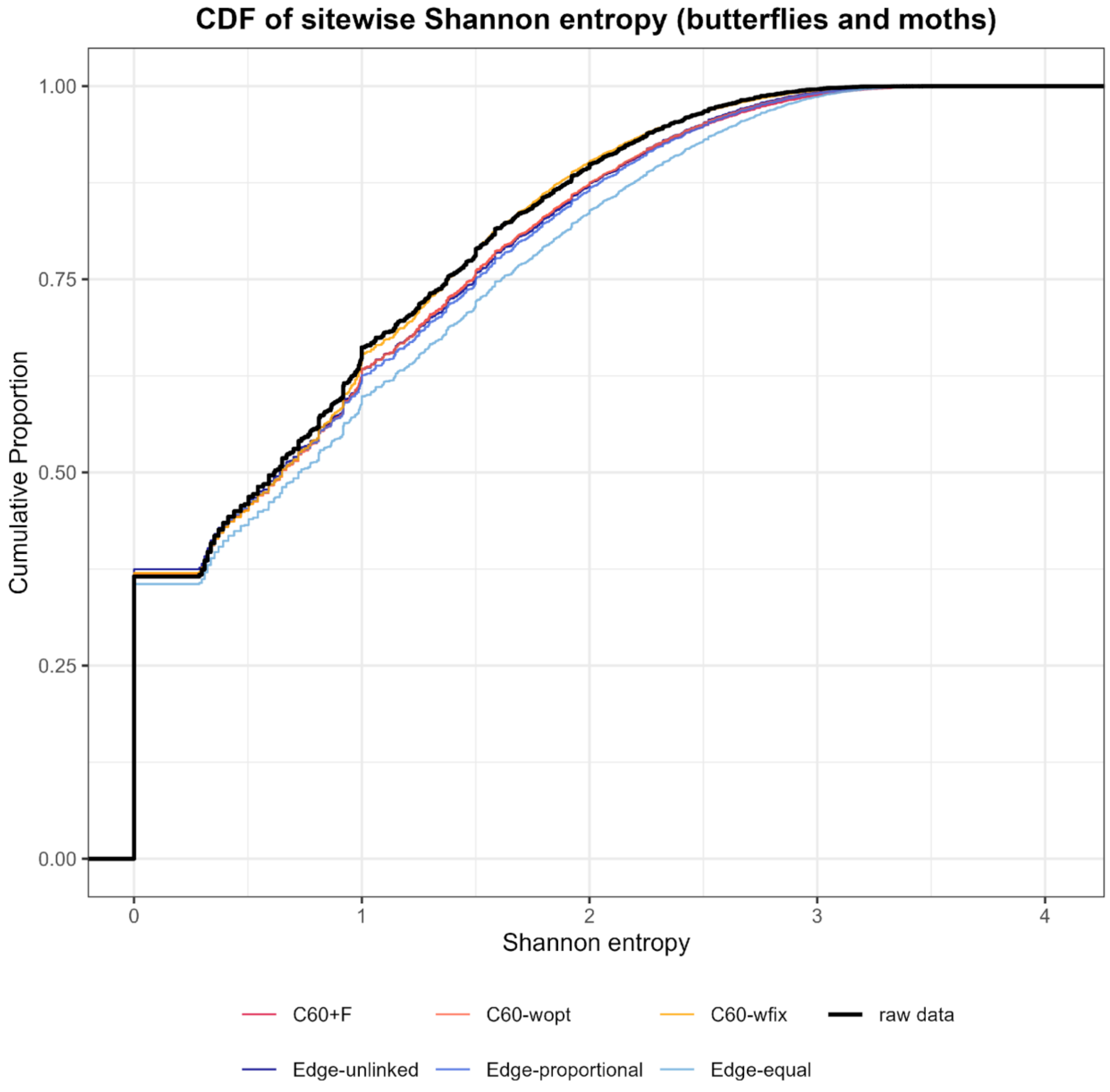

**Fig. S7.** Cumulative distribution function of site-wise Shannon entropy for 100 simulated datasets under six models compared to the raw birds dataset (subsampled to 20 taxa). For each of the six models, site-wise Shannon entropy values from 100 simulated replicate datasets were pooled into a single distribution, and the resulting CDF (coloured lines) is compared against the CDF of the empirical alignment (black line). Note that some model CDFs may closely overlap and appear visually indistinguishable in the plot.

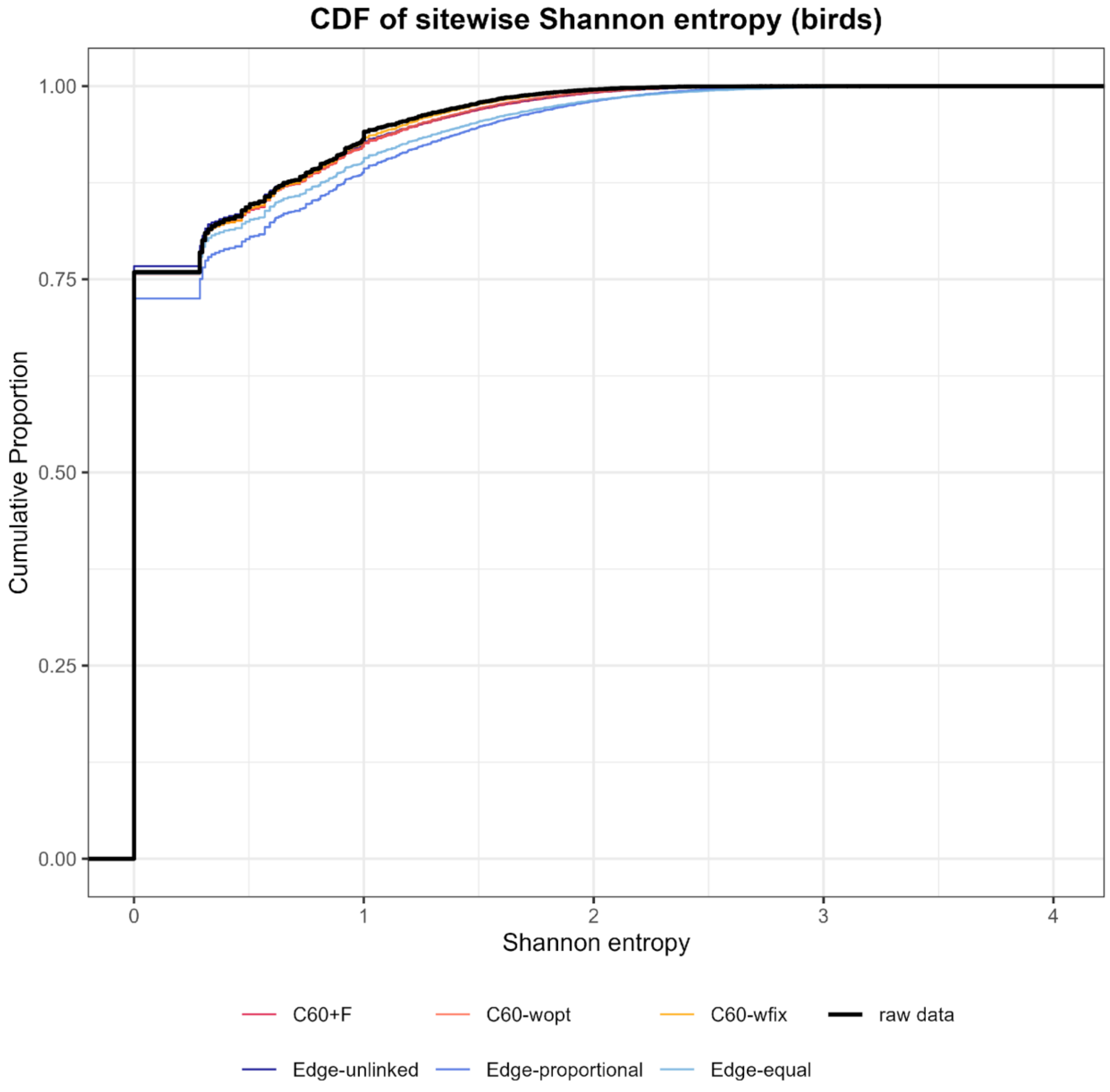

**Fig. S8.** Cumulative distribution function of site-wise Shannon entropy for 100 simulated datasets under six models compared to the raw green plants dataset (subsampled to 20 taxa). For each of the six models, site-wise Shannon entropy values from 100 simulated replicate datasets were pooled into a single distribution, and the resulting CDF (coloured lines) is compared against the CDF of the empirical alignment (black line). Note that some model CDFs may closely overlap and appear visually indistinguishable in the plot.

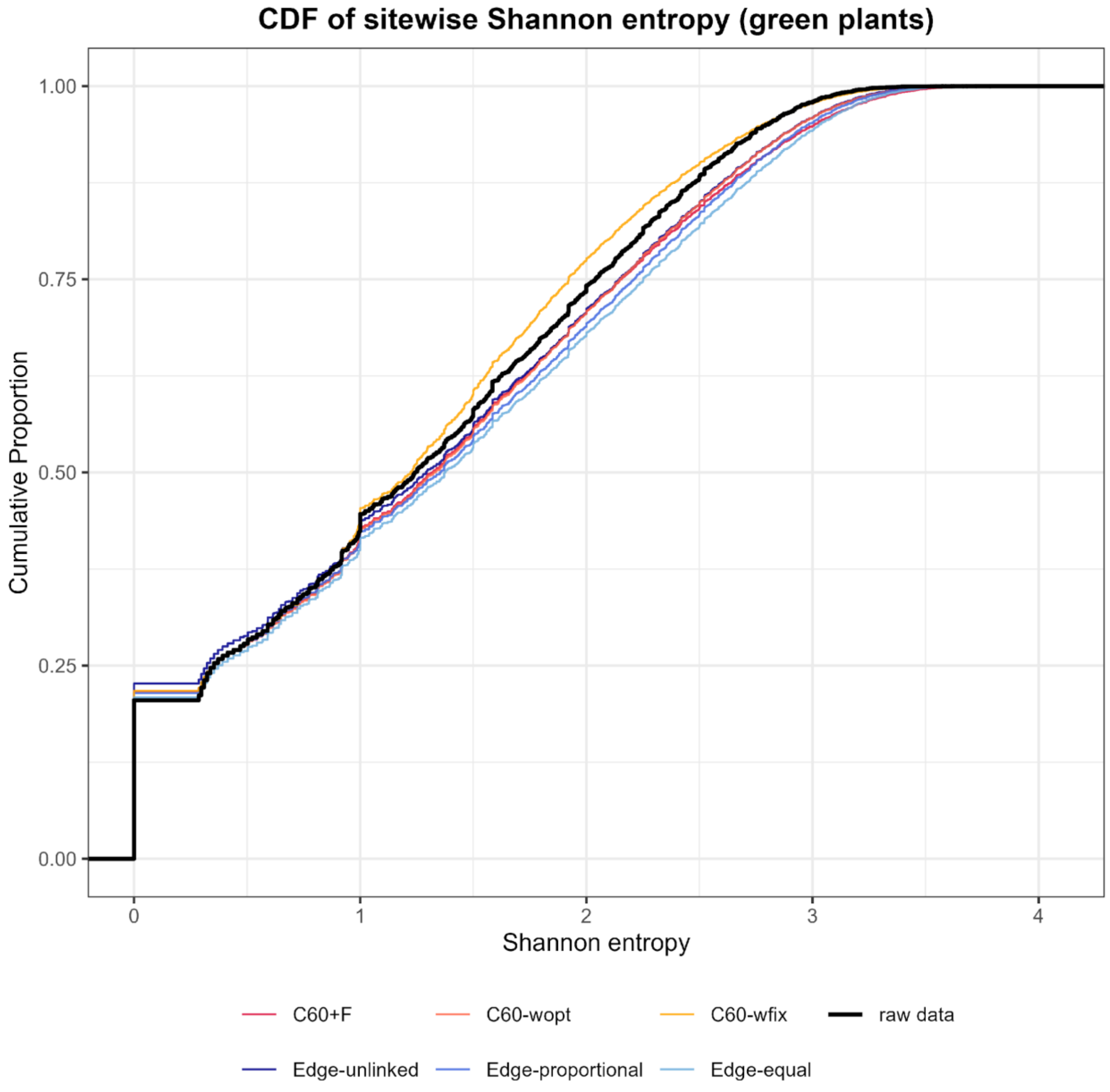

**Fig. S9.** Cumulative distribution function of site-wise Shannon entropy for 100 simulated datasets under six models compared to the raw legume dataset (subsampled to 20 taxa). For each of the six models, site-wise Shannon entropy values from 100 simulated replicate datasets were pooled into a single distribution, and the resulting CDF (coloured lines) is compared against the CDF of the empirical alignment (black line). Note that some model CDFs may closely overlap and appear visually indistinguishable in the plot.

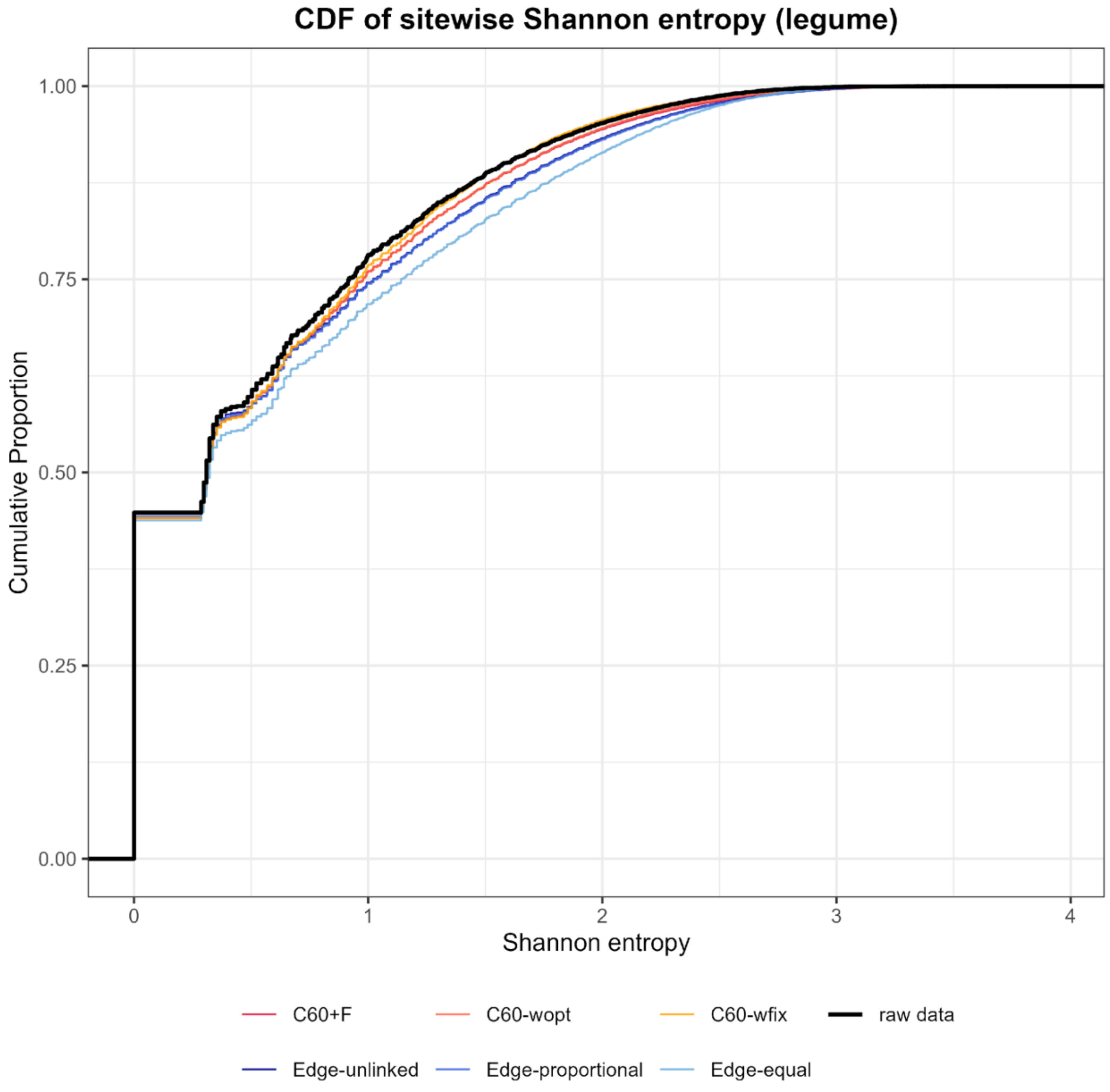

**Fig. S10.** Cumulative distribution function of site-wise Shannon entropy for 100 simulated datasets under six models compared to the raw Ascomycota dataset (subsampled to 20 taxa). For each of the six models, site-wise Shannon entropy values from 100 simulated replicate datasets were pooled into a single distribution, and the resulting CDF (coloured lines) is compared against the CDF of the empirical alignment (black line). Note that some model CDFs may closely overlap and appear visually indistinguishable in the plot.

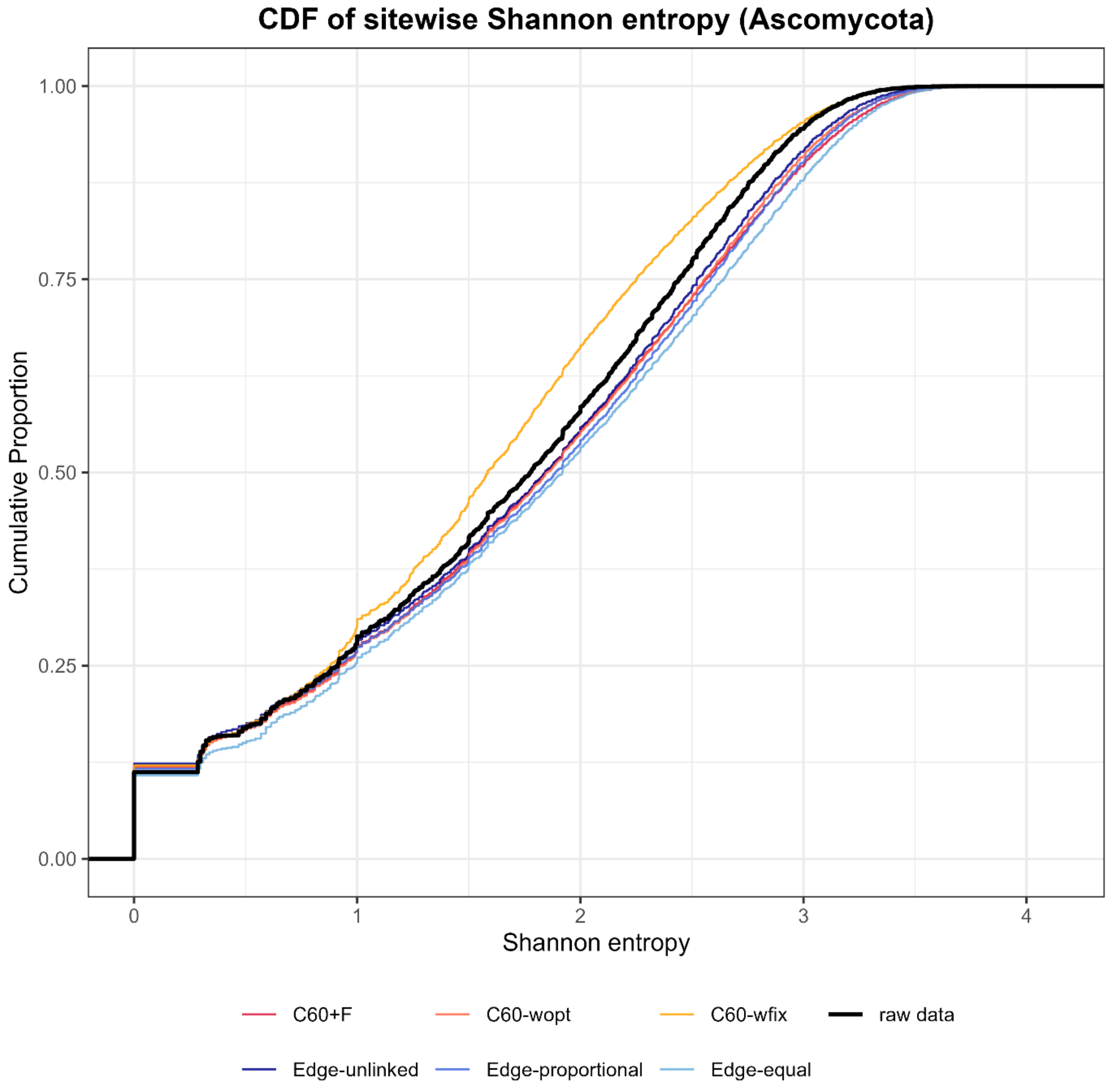

**Fig. S11.** Cumulative distribution function of site-wise Shannon entropy for 100 simulated datasets under six models compared to the raw budding yeast dataset (subsampled to 20 taxa). For each of the six models, site-wise Shannon entropy values from 100 simulated replicate datasets were pooled into a single distribution, and the resulting CDF (coloured lines) is compared against the CDF of the empirical alignment (black line). Note that some model CDFs may closely overlap and appear visually indistinguishable in the plot.

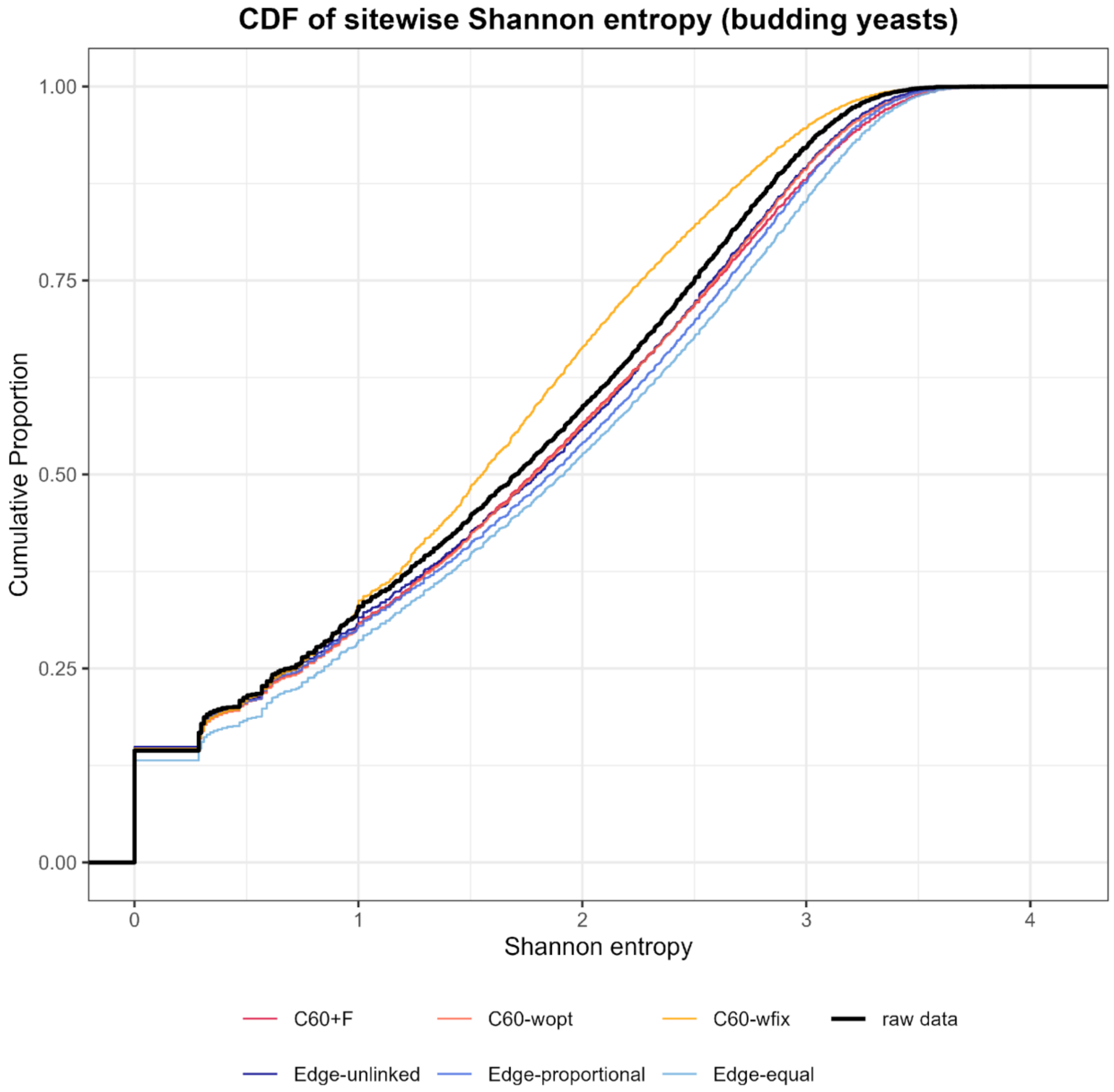

**Fig. S12.** Cumulative distribution function of site-wise Shannon entropy for 100 simulated datasets under six models compared to the raw cyanobacteria dataset (subsampled to 20 taxa). For each of the six models, site-wise Shannon entropy values from 100 simulated replicate datasets were pooled into a single distribution, and the resulting CDF (coloured lines) is compared against the CDF of the empirical alignment (black line). Note that some model CDFs may closely overlap and appear visually indistinguishable in the plot.

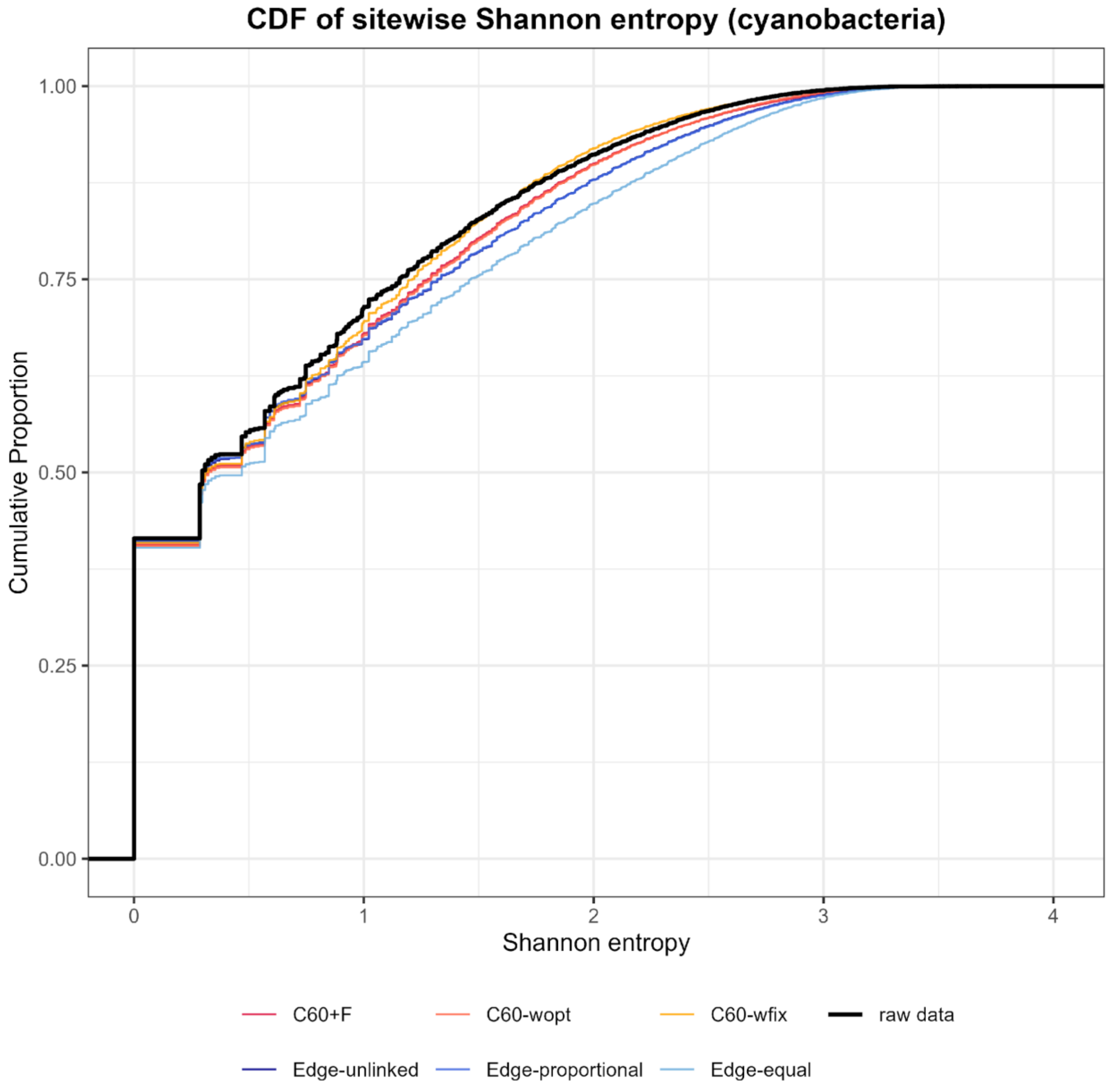

**Fig. S13.** Cumulative distribution function of site-wise Shannon entropy for 100 simulated datasets under six models compared to the raw archaea dataset (subsampled to 20 taxa). For each of the six models, site-wise Shannon entropy values from 100 simulated replicate datasets were pooled into a single distribution, and the resulting CDF (coloured lines) is compared against the CDF of the empirical alignment (black line). Note that some model CDFs may closely overlap and appear visually indistinguishable in the plot.

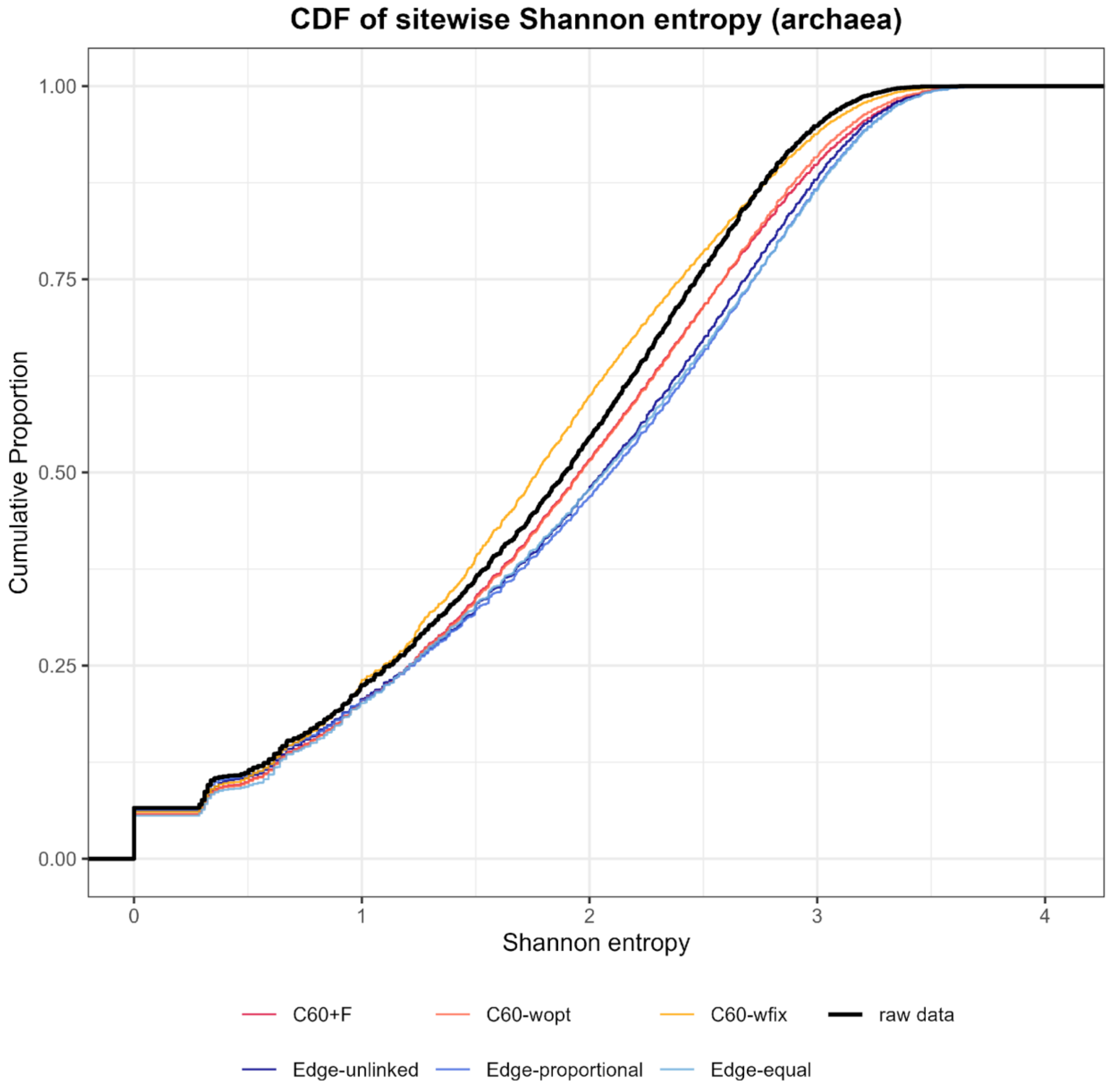

**Fig. S14.** All information scores of partition models on 20-taxon datasets. The x-axis shows the model types, and the y-axis shows the corresponding information criterion values. Numbers above the partition models indicate the optimal number of partitions inferred by PartitionFinder. Note that cAIC and AICc are expected to converge when the number of sites is much larger than the number of parameters, as the finite-sample correction term in AICc becomes negligible. This may cause their points to overlap and appear as a single point in some panels.

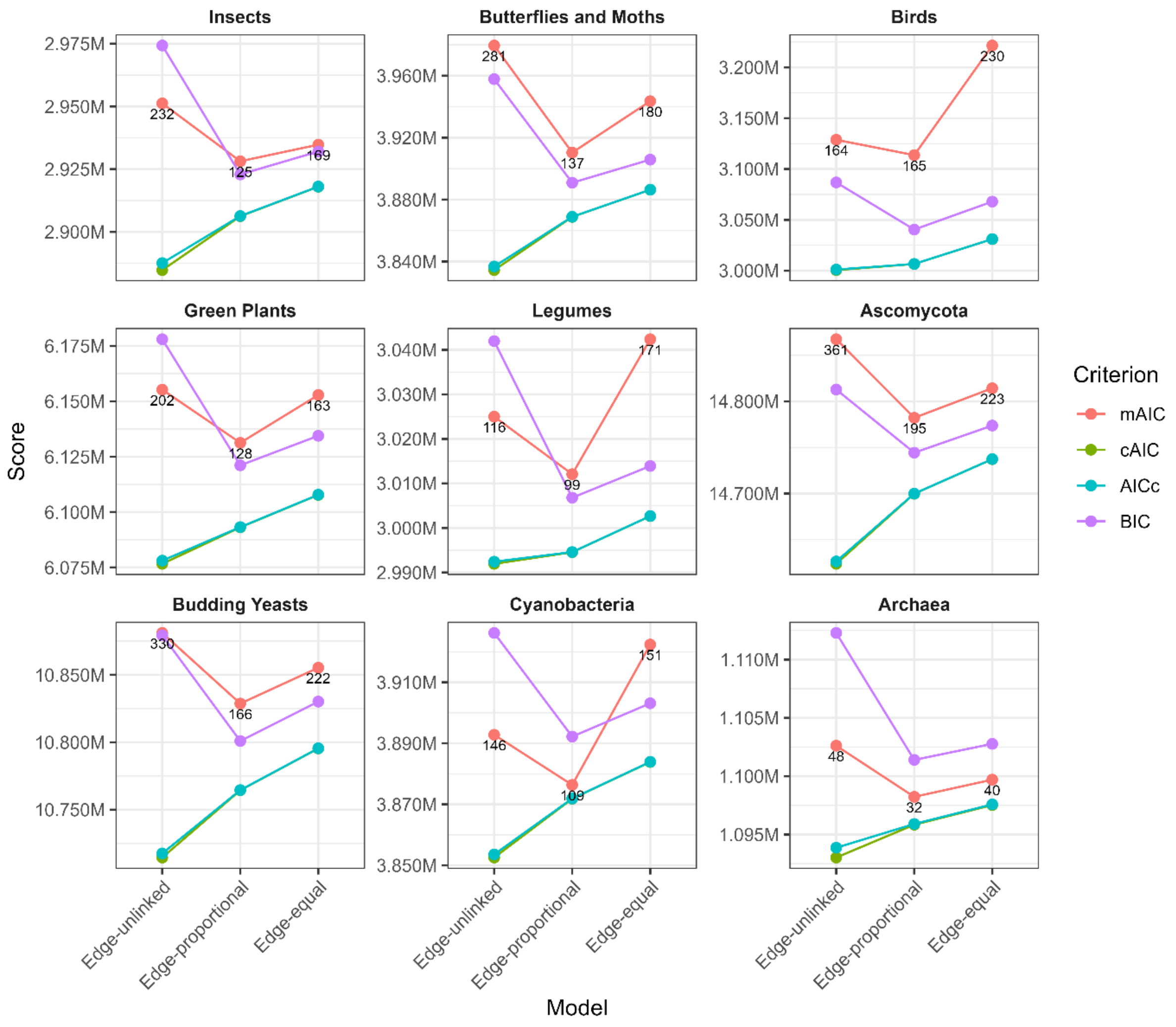

**Fig. S15.** All information scores of partition models on 10-taxon datasets. The x-axis shows the model types, and the y-axis shows the corresponding information criterion values. Numbers above the partition models indicate the optimal number of partitions inferred by PartitionFinder. Note that cAIC and AICc are expected to converge when the number of sites is much larger than the number of parameters, as the finite-sample correction term in AICc becomes negligible. This may cause their points to overlap and appear as a single point in some panels.

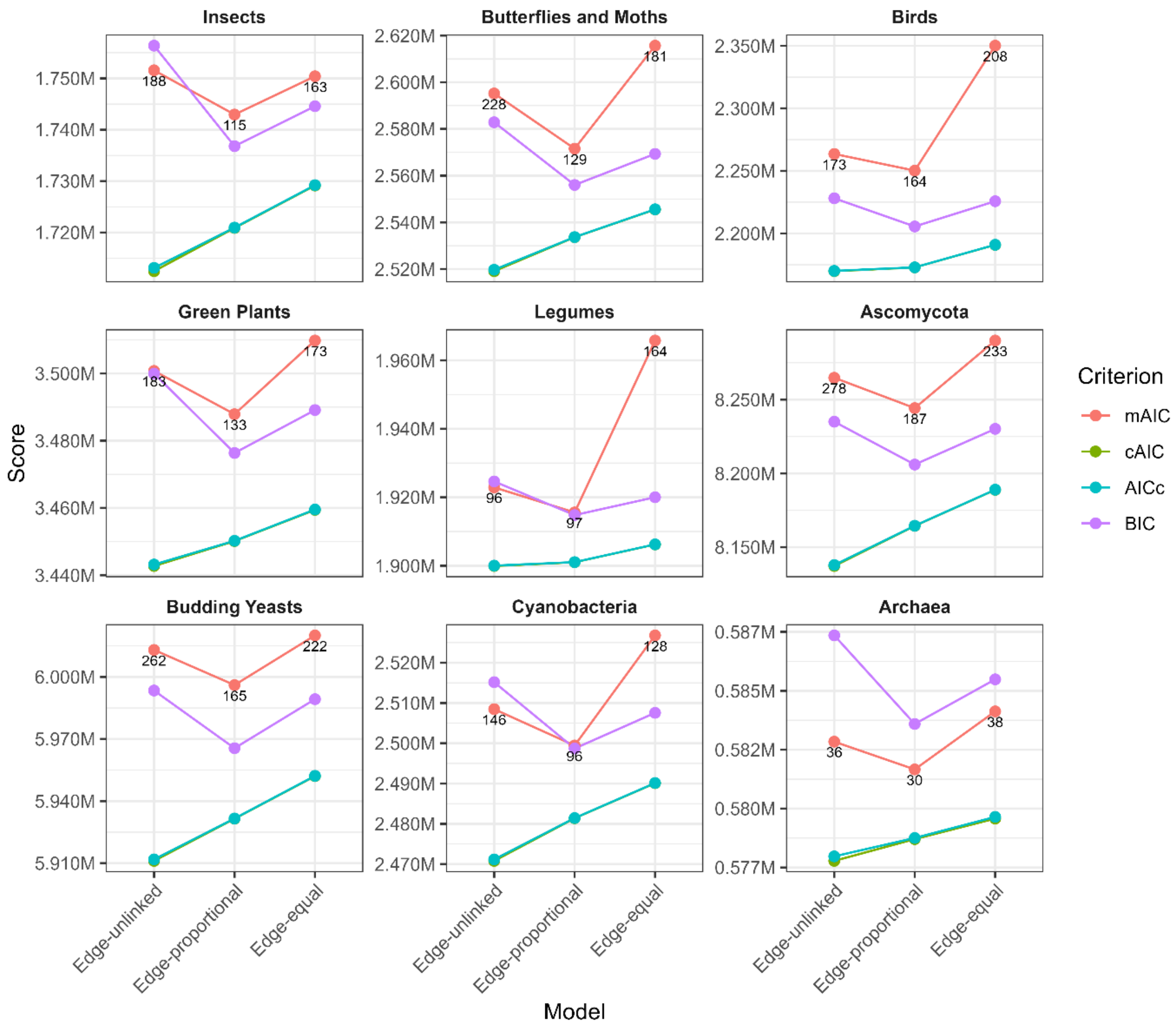

**Fig. S16.** All information scores of partition models on 5-taxon datasets. The x-axis shows the model types, and the y-axis shows the corresponding information criterion values. Numbers above the partition models indicate the optimal number of partitions inferred by PartitionFinder. Note that cAIC and AICc are expected to converge when the number of sites is much larger than the number of parameters, as the finite-sample correction term in AICc becomes negligible. This may cause their points to overlap and appear as a single point in some panels.

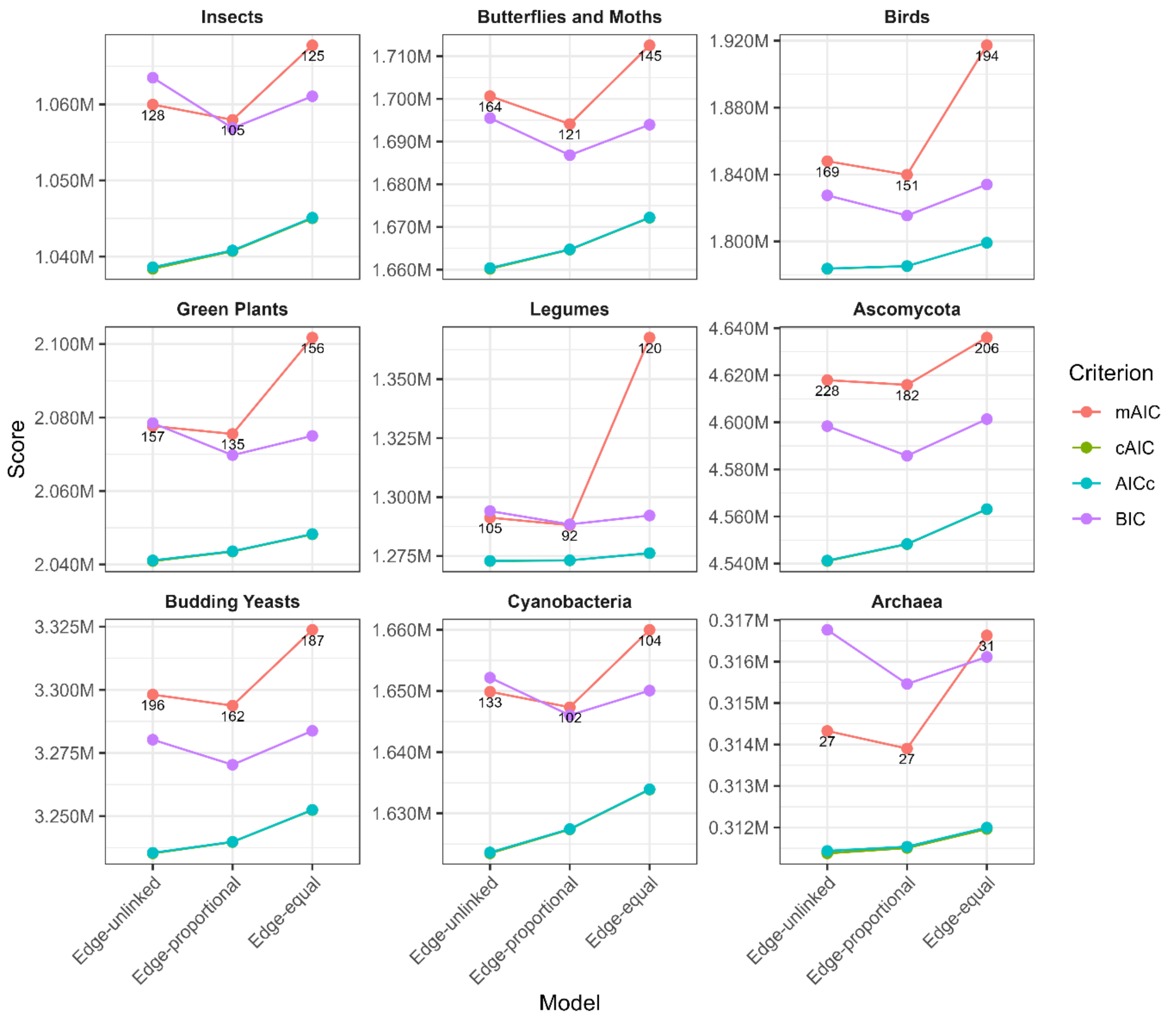

**Fig. S17.** All information scores of C60 model variants on 20-taxon datasets. The x-axis shows the model types, and the y-axis shows the corresponding information criterion values. Note that AIC and AICc are expected to converge when the number of sites is much larger than the number of parameters, as the finite-sample correction term in AICc becomes negligible. This may cause their points to overlap and appear as a single point in some panels.

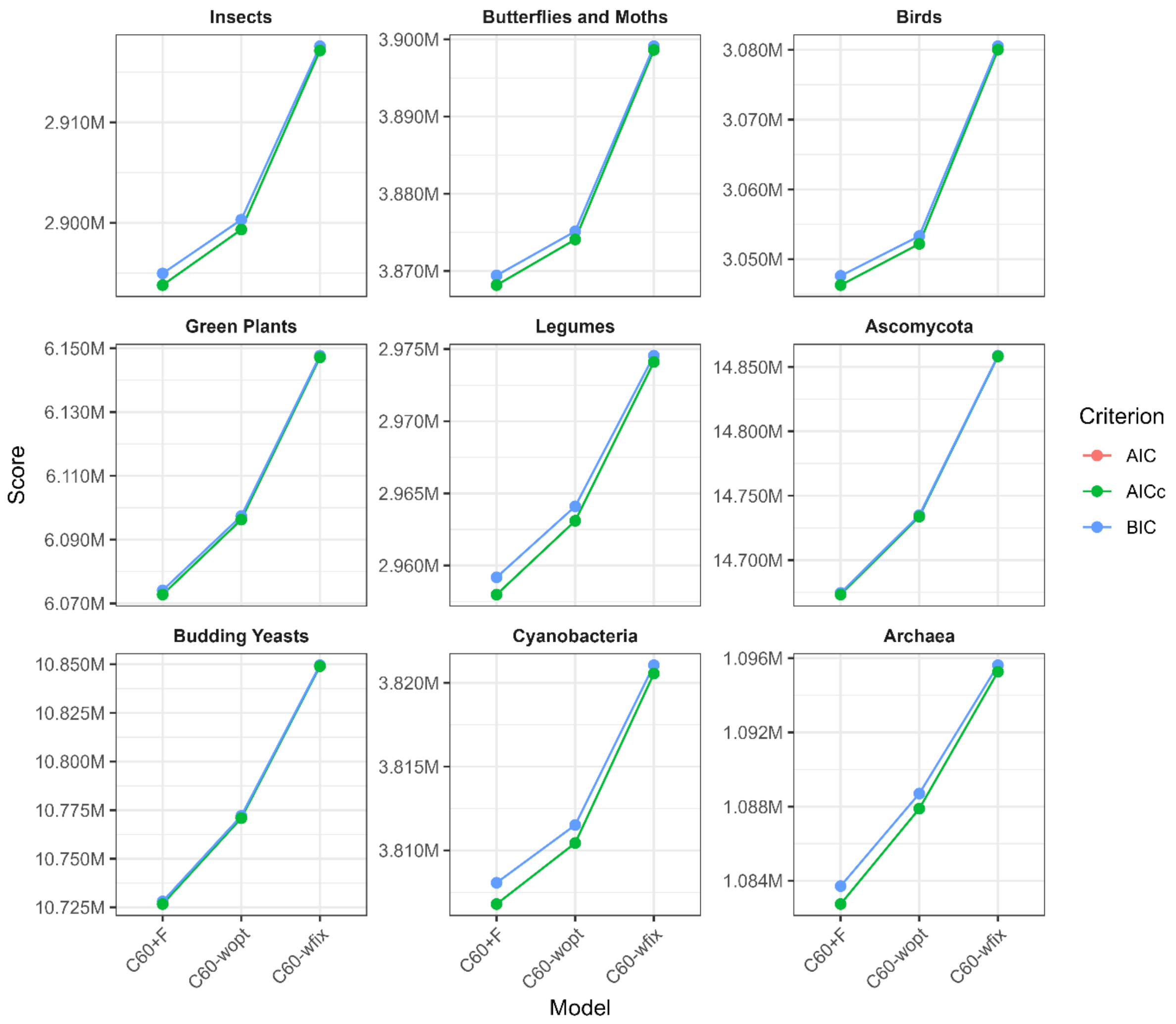

**Fig. S18.** All information scores of C60 model variants on 10-taxon datasets. The x-axis shows the model types, and the y-axis shows the corresponding information criterion values. Note that AIC and AICc are expected to converge when the number of sites is much larger than the number of parameters, as the finite-sample correction term in AICc becomes negligible. This may cause their points to overlap and appear as a single point in some panels.

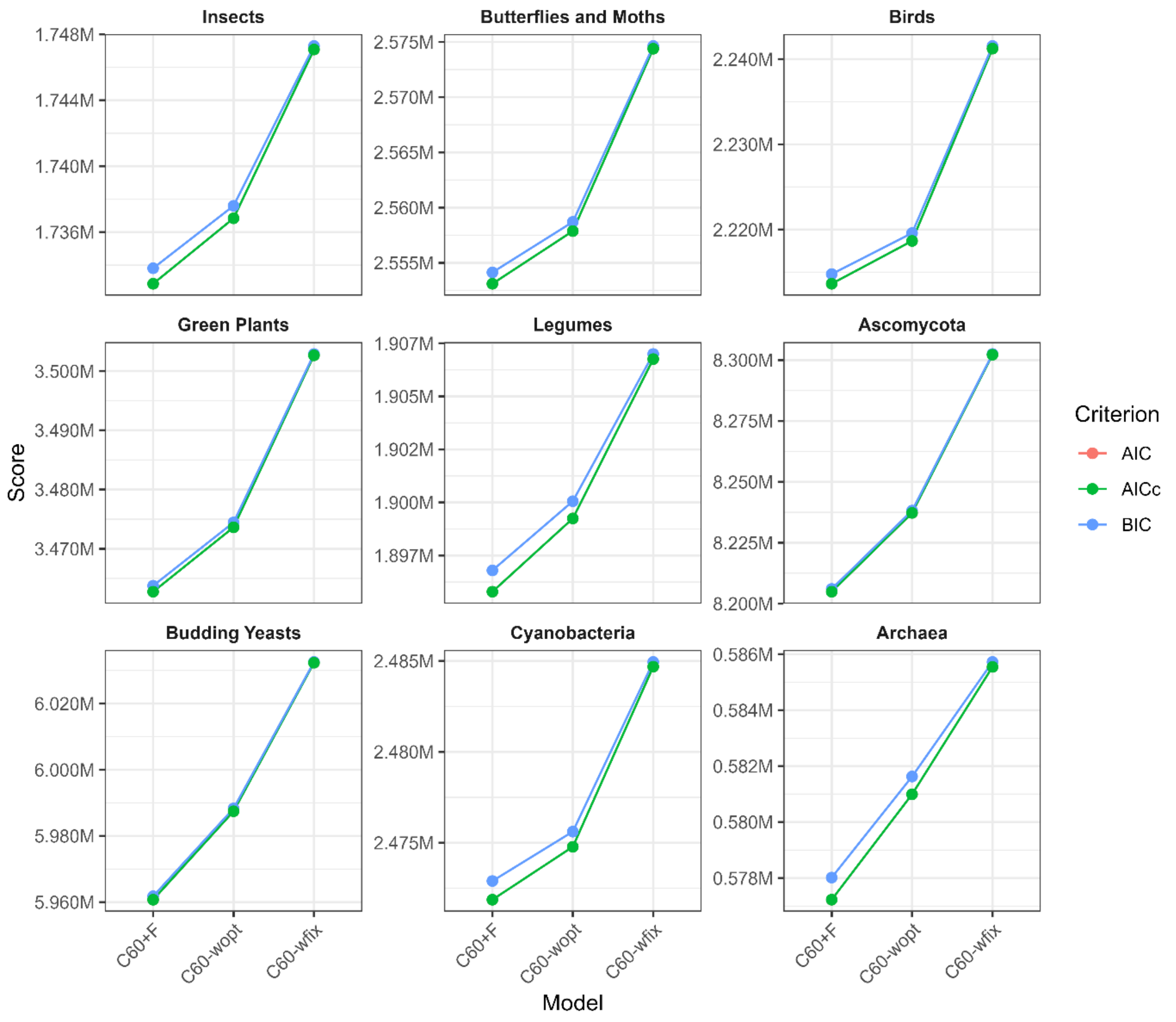

**Fig. S19.** All information scores of C60 model variants on 5-taxon datasets. The x-axis shows the model types, and the y-axis shows the corresponding information criterion values. Note that AIC and AICc are expected to converge when the number of sites is much larger than the number of parameters, as the finite-sample correction term in AICc becomes negligible. This may cause their points to overlap and appear as a single point in some panels.

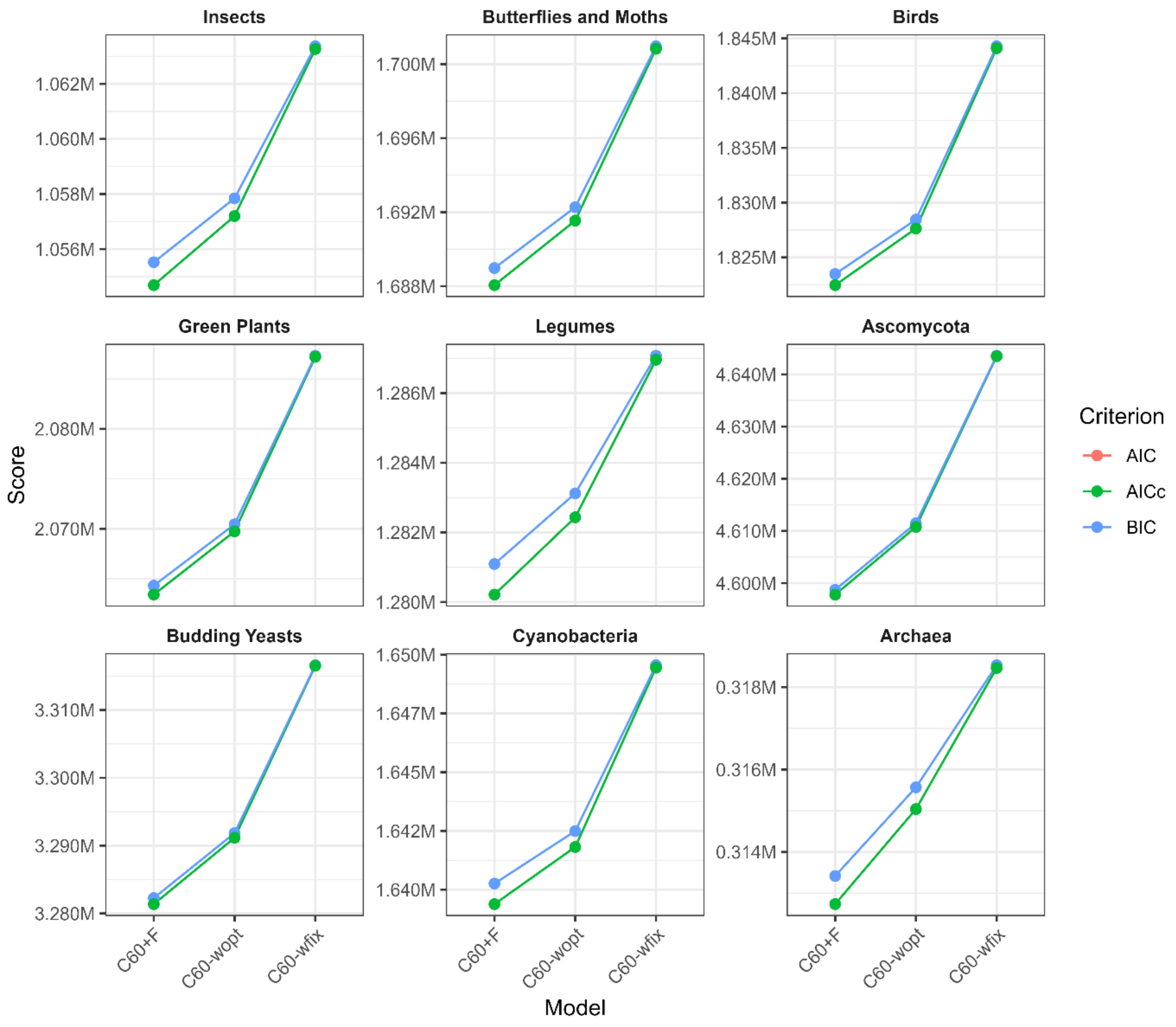

**Fig. S20.** AIC scores of C60, C10 and partition models on 20-taxon datasets. The x-axis shows the nine model types, and the y-axis shows the corresponding AIC values. Numbers above the partition models indicate the optimal number of partitions inferred by PartitionFinder.

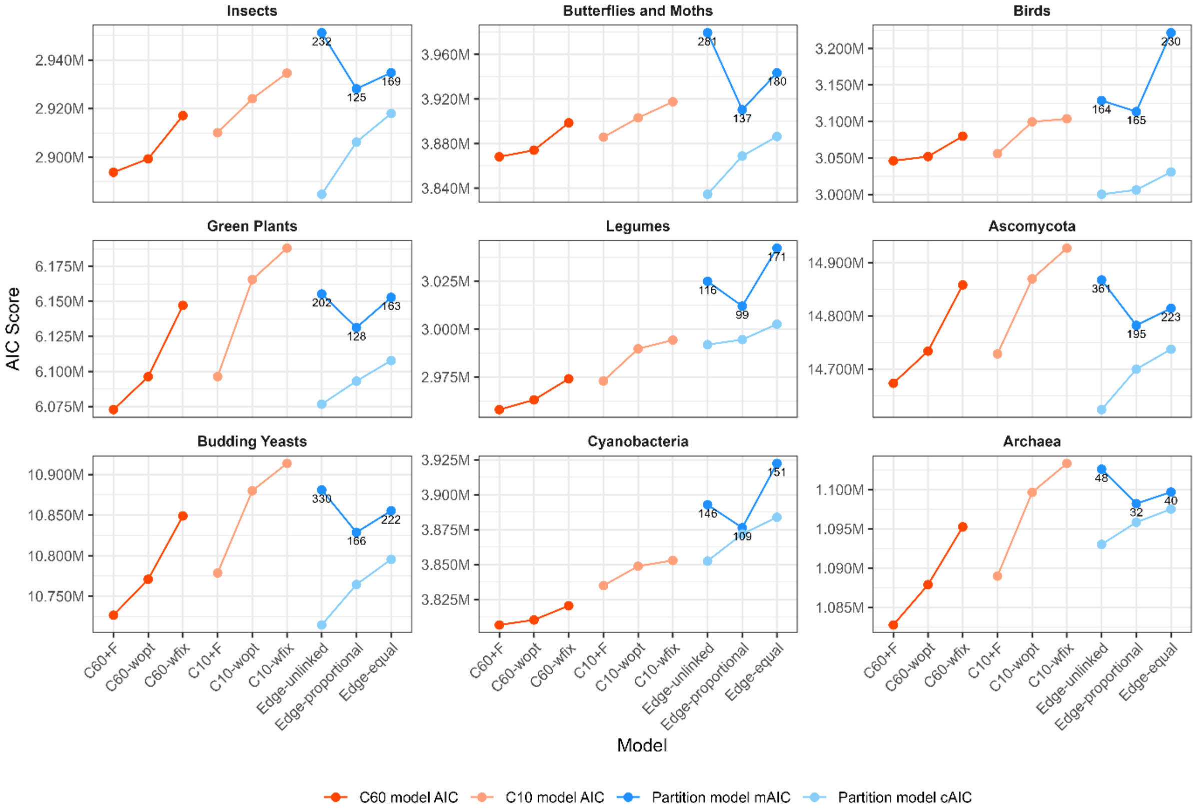
